# Molecular Characterization of Gene–Product Relationships in the Wheat Gluten Complex for Reduced-Immunogenicity Wheat Development

**DOI:** 10.64898/2026.10.12.758671

**Authors:** Gaganjeet, Gautam Saripalli, Zachary T. Jones, Anmol Kajla, Vijay Tiwari, Jacek Waga, Sachin Rustgi

## Abstract

Immune-mediated reactions to wheat gliadins, including celiac disease and wheat allergies, affect ∼1% of the U.S. population and are prevalent worldwide. Reducing the risk of inadvertent gluten exposure for individuals with celiac disease or wheat allergies requires the development of reduced immunogenicity wheat varieties which are suitable for human consumption. This study was designed with a view to improve our understanding of gene–protein relationships at the complex wheat *Gli1/Glu3* and *Gli2* loci, and to develop genetic and molecular resources for breeding reduced-immunogenicity wheat lines. We conducted comprehensive genomic, biochemical, and immunological analyses of diverse wheat genetic resources, including aneuploid stocks, chromosome deficiency and interstitial deletion lines, Wasko^ω−^, the landrace San Pastore, Pegaso near-isogenic lines lacking *Gli1/Glu3/Gli2* loci, and progenies derived from a Wasko^ω−^ × Pegaso^(6A–6D)α−^. Genomic characterization involved the targeted GlutEnSeq approach and PCR based DNA markers while biochemical and immunological analyses employed acid-urea PAGE, SDS-PAGE, and ELISA. Integration of these complementary approaches resolved gliadin gene–protein relationships at an unprecedented level. Notably, two F₂ lines exhibited substantial reductions (both α- and ω-gliadins) due to the successful stacking of parental mutations. The resulting F₂/F₃ progenies and associated DNA markers represent valuable resources for breeding wheat for reduced immunogenicity.

## 1. Introduction

Wheat (*Triticum aestivum* L.) is a cosmopolitan cereal crop that serves as a staple food for millions of people worldwide. Global wheat production reached 842.2 million metric tons in 2025/26, with China, India, Russia, and the United States ranking among the leading producers [1]. Nutritionally, wheat provides energy (∼72% carbohydrates by weight), protein (10–18% by weight), health-promoting dietary fibers (including arabinoxylans and β-glucans), and several bioactive compounds [2]. Wheat proteins are of particular importance because they confer the characteristic viscoelastic properties of wheat dough, and their composition in the grain determines its specific end uses. Broadly, wheat proteins are categorized into (i) gluten proteins (∼80% by weight), comprising approximately equal proportions of gliadins and glutenins, and (ii) non-gluten proteins, which consist of albumins and globulins [3]. Among the seed storage proteins that accumulate in starchy endosperm cells during grain development, gluten proteins are the most complex and are encoded by 12 loci distributed across two groups of three homoeologous chromosomes [4]. Within the gluten protein complex, gliadins are classified into sulfur-rich α- and γ-gliadins and sulfur-poor ω-gliadins, whereas glutenins are large polymers composed of low- and high-molecular-weight subunits linked by disulfide bonds [4]. The α-gliadin genes (27 in ‘Chinese Spring’ [CS] and 44 in ‘JM44’) are located at the *Gli-2* loci on the short arms of the group 6 chromosomes. In contrast, the γ-gliadin (12 in CS and 17 in JM44), ω-gliadin (6 in CS and 9 in JM44), and low-molecular-weight glutenin subunit (LMW-GS) genes (15 in CS and 21 in JM44) are located at the *Gli-1* and *Glu-3* loci on the short arms of the group 1 chromosomes [5–8].

In addition to having wide implications for determining end-use performance and nutritional quality, gluten proteins are also implicated in triggering immune reactions in approximately 5% of the global population [9]. Among gluten-induced immune reactions, celiac disease (CD) is one of the most common chronic, immune-mediated enteropathic conditions [10]. Other, less common reactions involve allergic responses to wheat proteins, including life-threatening exercise-induced anaphylaxis. In addition to these conditions, many individuals experience non-allergic, non-celiac, and extraintestinal symptoms upon wheat exposure, collectively known as non-celiac wheat sensitivity (NCWS). Currently, the most effective treatment for these conditions is strict avoidance of wheat and other gluten-containing cereals, or limiting gluten content in food items to an internationally approved 20 ppm threshold [11]. However, accidental gluten exposure is common [12]. In addition, long-term adherence to a wheat-free (gluten-free) diet has been associated with adverse health consequences [13–14]. Therefore, efforts are being made to develop various therapies and food alternatives, such as vaccines, oral enzyme therapies, and reduced-immunogenicity wheat genotypes [15–19]. However, oral enzyme therapies tested and released in certain parts of the world, as well as experimental vaccines such as Nexvax2 [20], have proven ineffective. Consequently, developing reduced-immunogenicity wheat lines appears to be a more practical alternative.

Different approaches have been attempted to develop reduced-immunogenicity wheat lines, including (i) RNA interference, (ii) CRISPR-mediated gene editing, and (iii) crossbreeding. A sizable amount of success has been achieved using the first approach, where a significant reduction in the content of immunogenic gliadins was observed. These reduced-gliadin lines (specifically, E82) produced acceptable-quality bread, exhibited a lower ability to stimulate T-cell immune responses, and generated non-significant immune activation upon consumption compared to wild-type wheat [21]. Similarly, CRISPR/Cas9- and CRISPR/Cas12a-based genome editing has enabled the development of lines with significant reductions in immunogenic gliadins [15,16]. These genotypes have been tested using various biochemical and immunological methods and appear promising; however, they require further improvement. Attempts to enhance these lines included stacking traits from different methods through crossbreeding, but only limited improvement over the RNAi line was achieved [16]. There have also been proposals to use homologous recombination to edit specific epitopes by combining the delivery of gene-editing reagents with an oligonucleotide to precisely modify epitope sequences. However, this approach has only been applied to a limited number of well-characterized epitopes [22], and multiple edits would need to be stacked within a single line to be effective. Furthermore, because any undigestible peptide of approximately nine amino acids can be a potential epitope—and given that epitope characterization is still ongoing—it is difficult to achieve substantial success with this strategy.

Transgenic approaches suffer from limited public acceptance and face regulatory hurdles due to the presence of foreign DNA. In contrast, gene-edited materials carry site-specific mutations at target sites and have been shown to lack foreign DNA [16]. However, these methods often involve biomolecule delivery techniques that overlap with conventional transgenesis, placing them within regulatory debates and complicating approval processes. Despite recent developments in which governments around the world have approved the release of such materials, questions remain regarding public acceptance. This issue was addressed in a recent study [23], which found that public opinion varies by crop with respect to the acceptance of gene-edited materials. The study suggested that while gene editing in wheat is considered morally acceptable, there is little intention among consumers to purchase and consume gene-edited reduced-gluten products [23]. In contrast, respondents expressed a greater willingness to accept similar materials developed through conventional crossbreeding.

This highlights the need for conventional breeding approaches to develop wheat cultivars that are widely acceptable. In the past, this approach has been successful in stacking three homoeologous ω-gliadin deletion mutants [24], two α-gliadin null mutations [25,26], and two ω-gliadin and two α-gliadin mutations [27]. However, although these genotypes stacked mutations in two to four gliadin genes, they still retained a significant amount of immunogenic gluten proteins. Additionally, the genomic locations of gluten genes have been known for some time, but a connection between the genes and their products has not yet been established. Therefore, the present study is designed to fill this gap with two major objectives: (i) to establish a connection between gliadin genes and their products through detailed evaluation of wheat aneuploid stocks and lines carrying terminal deficiencies and interstitial deletions, and (ii) to develop genetic resources (germplasm lines with stacked deletions in up to five gliadin genes) and genomic resources (DNA markers) to breed wheat cultivars suitable for consumption by wheat allergic individuals and celiac patients. Overall, this study will enhance our understanding of the gluten gene–product relationship and will develop genetic and genomic resources for breeding wheat with reduced levels of immunogenic gluten proteins.

## 2. Materials and Methods

### 2.1. Plant material

A wide range of wheat genetic stocks were used in this study (Table S1). Briefly, these genotypes include nullisomic–tetrasomic (NT) lines for the chromosome groups 1 to 7, deletion lines for group 1 and group 6 chromosomes in the CS background; lines carrying single mutations at the *Gli-A1* (Pegaso^−1A^), *Gli-B1* (Pegaso^−1B^), and *Gli-D1* (Pegaso^−1D^) loci; as well as double mutations at *Gli-A1/D1* (Pegaso^−1A/1D^), *GliB1/D1* (Pegaso^−1B/1D^), and *Gli-A2/D2* (Pegaso^(6A–6D)α-^) loci in the ‘Pegaso’ background. In addition, a triple mutant carrying mutations in *Gli-A1*, *Gli-B1*, and *Gli-D1* (Wasko^ω-^), gamma radiation–induced mutants in the background of a hard red winter wheat cultivar ‘Jagger,’ and a wheat landrace, ‘San Pastore,’ were also included. These wheat lines were used for physical mapping of gluten genes and their products through various methods (see below). The stacked mutants Wasko^ω-^ and Pegaso^(6A–6D)α-^ were further combined via crossing, and the F₂ and F₃ generations were characterized for their gluten profiles and evaluated for their genetic makeup. Seeds of the different wheat lines were grown in 9-inch pots. At the two-leaf stage, plants were transferred to a Conviron growth chamber (Model A1000, Controlled Environments Ltd., Winnipeg, MB, Canada) set at 4–8 °C for six weeks (8 h photoperiod) to induce flowering. Following vernalization, plants were transferred to the greenhouse facility at the Pee Dee Research and Education Center, Florence, SC, for crossing.

### 2.2. Extraction and profiling of gliadins and glutenins

Seeds of each genotype were divided into endosperm and embryo halves prior to gliadin and glutenin extraction using a 3D-printed handheld device. The endosperm half was used for the sequential extraction of gliadins and glutenins following [28], while the embryo half of selected lines was used to advance plants to the next generation, as described earlier. Briefly, gluten extraction involved the following steps: (i) The endosperm half of the seed was ground into a fine flour. (ii) A total of 210 µL of 60% ethanol (v/v) was added to the flour, vortexed for 45 s to homogenize the mixture, shaken for 1 h at 200 rpm in an orbital shaker (at room temperature), and centrifuged at 10,000 × g for 20 minutes to extract gliadins. (iii) After transferring the supernatant (containing gliadins) to a 1.5 mL tube, the pellet was re-extracted for glutenins by adding freshly prepared glutenin extraction buffer containing 60% 1-propanol (v/v), 2% dithiothreitol (DTT) (w/v), 2 M urea (w/v), and 0.05 M Tris-HCl. (iv) The pellet was vortexed until completely dissolved in the buffer and then shaken at 200 rpm in an incubator shaker at 60 °C for 2 h. Subsequently, the mixture was centrifuged at 8,000 rpm for 20 min, and the supernatant containing glutenins was collected in a separate tube. Following extraction, both gliadins and glutenins were stored at 4 °C until further use. In total, 10 seeds from each genotype were analyzed separately to characterize gliadin and glutenin compositions using acid-urea PAGE and SDS-PAGE, respectively, following standardized protocols described by [29] and [30].

The results of gliadin and glutenin profiling are presented based on the presence or absence of specific gliadin and glutenin bands, as well as differences in band intensities. Protein reduction was quantified as a percentage based on band intensity relative of wild-type Pegaso in densitograms generated from protein gel images using Image J [30].

### 2.3. Statistical analysis

The densitometric estimates obtained from protein gels were used to perform analysis of variance (ANOVA) among the eight genotypes listed in Table S1, including Pegaso wild type, Pegaso near-isogenic lines (NILs), and Wasko^ω⁻^ (*Gli-A1*, *Gli-B1*, and *Gli-D1* triple null). For this analysis, protein profiles from three technical replicates of each genotype were included, and a one-way ANOVA was conducted using R. Pairwise comparisons among genotype means were performed using Tukey’s honestly significant difference (HSD) post hoc test implemented in the Agricolae package in R [31].

### 2.4. Gliadin quantification using sandwich R5 ELISA

A set of 14 selected samples (Table S1) was analyzed for total gliadin content using the R5 antibody (Neogen Veratox® Gliadin R5 ELISA kit; Neogen, Lansing, MI, USA). These samples included the wheat reference genotype, CS; two mutant lines used for crossing (Wasko^ω⁻^ and Pegaso^(6A–6D)α⁻^); eight F₂ lines; and three F₃ lines. Samples for ELISA were prepared according to the manufacturer’s instructions using 10 mM phosphate-buffered saline (PBS). Gliadins were extracted as described above and quantified using a Qubit 4 Fluorometer (Thermo Fisher Scientific, Massachusetts, USA). The extracted gliadins were diluted in 1% PBS at a 1:6000 ratio, and the assay was performed in duplicate. Subsequently, the optical density (OD) values obtained for the samples were compared with those of the standards and adjusted for the dilution factor. The results were expressed as a percentage of the parts-per-million (ppm) value obtained for CS.

### 2.5. GlutEnSeq analysis

A subset of 12 lines (Table S1), selected based on their protein profiles, was used for detailed characterization using a next-generation selective sequencing (NGS) approach, GlutEnSeq [32]. The GlutEnSeq results were validated using PCR-based markers. For this purpose, 100 mg of wheat leaf tissue was collected and lyophilized for 72 h. Genomic DNA was extracted using the E.Z.N.A.® Plant & Fungal DNA Extraction Kit (Omega Bio-tek Inc., Georgia, USA), following the manufacturer’s instructions. DNA was quantified using a NanoDrop spectrophotometer (Thermo Fisher Scientific, Massachusetts, USA) and the concentration was adjusted to 30 ng/µL. Samples were then dispatched for GlutEnSeq to Daicel Arbor Biosciences, USA. Briefly, the following steps were involved in library preparation and NGS analysis: (i) A single capture pool was prepared for each DNA sample (up to 1000 ng DNA). (ii) Captures were performed following the myBaits v5.03 protocol using a custom myBaits bait set (design ID: D10749GLTNS), with overnight hybridization and washes at 65 °C. Up to 80% of the gDNA was filtered out using this bead-based method, and the remainder was prepared into a dual-indexed, Illumina-compatible library with an average insert size of approximately 500 bp. The indexed libraries were quantified using a spectrofluorometric assay. The capture pool was dried down to 7 µL by vacuum centrifugation. Post-capture, the reactions were amplified for 10 cycles, quantified again using a spectrofluorometric assay, and visualized using the TapeStation 4200 (Agilent) platform with High Sensitivity D5000 tape. The pool was sequenced on the Element Biosciences AVITI system using a partial PE150 high-throughput flow cell to approximately 1 Gb per library. Demultiplexed FASTQ data were used for downstream bioinformatics analysis.

### 2.6. Bioinformatics analysis

Paired-end GlutEnSeq reads were aligned against the CS wheat reference genome (RefSeq v2.1) and the bait sequences for gliadin genes [32] using BWA v0.7.19 [33]. The resulting ‘.sam’ alignment files were converted to ‘.bam’ format using SAMtools. To generate coverage data for genomic regions encompassing the gliadin genes, ‘.bed’ files were first created using gliadin gene coordinates from [34]. These ‘.bed’ files, together with the ‘.bam’ alignment files, were used as input in BEDTools to calculate coverage density values. The reads were normalized against the housekeeping gene *TaPFT1* following [32]. Coverage density values for different genotypes across various gluten gene sequences were calculated relative to the wild-type Pegaso. This coverage data was used to generate coverage plots (with gene order determined from the CS reference sequence) using Microsoft Excel. Deletion sizes in the Pegaso NILs, Pegaso^−1A^, Pegaso^−1B^, Pegaso^−1D^, Pegaso^−1A/1D^, Pegaso^−1B/1D^, and Pegaso^(6A-6D)α-^, and Wasko^ω⁻^ (*Gli-A1*, *Gli-B1*, and *GliD1* triple null), were interpreted by comparing their coverage values with those of the wild-type Pegaso (Pegaso WT). Similarly, deletions in the F_2_ and F_3_ progenies of the Wasko^ω⁻^ × Pegaso^(6A–6D)α⁻^ cross were interpreted based on comparisons with the parental genotypes. The results were visualized as coverage plots and heatmaps using Microsoft Excel and ClustVis (https://biit.cs.ut.ee/clustvis/<u>; accessed on 15th April,</u> <u>2026),</u> respectively [35]. In addition, gliadin banding patterns of F_2_ population were scored as present (+), absent (-) or heterozygous (0.5) (based on band intensity) and were coded as 1 (blue), 0 (light yellow), and 0.5 (orange), respectively. The resulting genotype by band matrix was visualized in v4.3.2. using. circlize v0.4.16 [36] and ComplexHeatmap v2.18.0 packages [37].

### 2.7. Molecular marker analysis

The deletions identified through GlutEnSeq analysis were further validated using PCR-based markers. In total, 30 markers were used, of which 4 were developed in this study, while the remaining markers were obtained from the literature (Table S2). PCR amplification was performed in a 20 μL reaction volume containing 1 U *Taq* DNA polymerase (MyTaq, Meridian Life Science Inc., USA), 1× *Taq* buffer with 2.5 mM dNTPs, and 10 pmol each of forward and reverse primers (MilliporeSigma, St. Louis, USA). Wheat cultivars CS and Pegaso were used as positive controls for each marker. Additionally, each marker was tested for sub genome specificity using DNA extracted from CS nullisomic–tetrasomic lines for group 1 and group 6 chromosomes. A housekeeping wheat gene, *Actin*, was used as a loading control [38]. The presence or absence of bands generated by different DNA markers across wheat genotypes was compared with the GlutEnSeq results and used for validation.

### 2.8. Epitope analysis

*In silico* analysis of immunogenic epitopes was conducted using GlutEnSeq reads for all the 12 samples. Specifically, the sequences were examined for reductions in the number of hits against immunogenic gliadin peptides. For this purpose, GlutEnSeq reads were subjected to BLASTx analysis against a locally prepared database of CD epitopes [39]. The GlutEnSeq reads in “.fasta” format was used as query in the local BLAST search against the epitope database. A total of 19 epitope sequences were used for the BLAST analysis, including 2 epitopes for ω-gliadins, 11 for γ-gliadins, and 6 for α-gliadins. The BLASTx option was employed to compare gliadin protein sequences against all possible reading frames of CS protein sequences. The following command was used for local BLAST analysis (e.g., GG01L6: epitopegroup6): “blastx -query GG01L6fasta.fasta -db epitopegroup6db -out epitopegroup6L6.txt -outfmt 6 -evalue 1000 - word_size 2 -matrix PAM30 -gapopen 9 -gapextend 1 -max_target_seqs 10.” Only BLASTx hits with up to two mismatches and a length of 9 amino acids were considered for interpretation. A high percent identity threshold was maintained to minimize misinterpretation. Furthermore, the abundance of epitopes corresponding to ω-, γ-, and α-gliadins was determined relative to the control. Bar graphs were generated using the percentage values of epitopes for ω-, γ-, and α-gliadins across each sub-genome. For all comparisons, Pegaso-WT was used as the control (set at 100% epitope frequency), and the abundance of epitopes in other samples was calculated relative to this reference.

### 2.9. Estimation of various seed parameters

Various seed parameters for all mutants, Pegaso-WT, and one F_3_ progeny were estimated using SmartGrain software ver. 1.2 [40]. The parameters measured included seed area, perimeter, length, width, length-to-width ratio, and circularity. The 100-seed weight of all genotypes was also estimated using the weight of 50 seeds for the mutants and Pegaso-WT, and 10 seeds for the crossed F_3_ progeny (Wx P-P8-7-9_F3).

## 3. Results

### 3.1. Development of the F_2_ population and its biochemical characterization

A total of 71 F_1_ seeds were obtained from a cross between Wasko^ω−^ and Pegaso^(6A–6D)α−^, including 18 seeds from a reciprocal cross. These 71 seeds produced 53 F_2_ plants; remining 18 seeds either did not germinate or plants died prematurely. From each plant, 10 seeds were evaluated for protein content and gluten composition. The F₂ population, parental genotypes, and other wheat genotypes listed in (Table S1) were profiled for gliadin and glutenin composition using acid-urea and SDS–polyacrylamide gel electrophoresis. In addition, a subset of 14 selected genotypes was evaluated for gliadin content using ELISA (see Materials and Methods for details).

### 3.2. Gliadin profiling of wheat genotypes

Gliadin profiling of different wheat genotypes led to several important observations. Given the diversity of materials studied and the objectives they serve, the results were divided into subsections.

(i) *Parental genotypes and F₂ lines*: Both parental genotypes, Pegaso^6A-6Dα⁻^ and Wasko^ω⁻^, exhibited the expected banding patterns. Pegaso^6A-6Dα⁻^ showed an absence of α-gliadin bands assigned to wheat chromosomes 6A and 6D, resulting in an approximately 80% reduction in α-gliadins relative to Pegaso-WT. Wasko^ω⁻^ lacked ω-gliadin bands associated with group 1 chromosomes. In addition to ω-gliadin deficiency, Wasko^ω⁻^ also exhibited an approximately 30% reduction in γ-gliadins estimated based on the densitometric analysis of protein gel images (Fig. 1A). Analysis of other group 1 chromosome mutants in the Pegaso background—including Pegaso^−1A^, Pegaso^−1B^, Pegaso^−1D^, and double-null mutants including Pegaso^−1A/1D^ and Pegaso^−1B/1D^—revealed the absence of gliadin bands assigned to specific sub genomes (Table S3; Fig. 1B). The densitometric estimates for the eight wheat genotypes, including Pegaso-WT, Pegaso NILs (Pegaso^−1A^, Pegaso^−1B^, Pegaso^−1D^, Pegaso^−1A/1D^, Pegaso^−1B/1D^, and Pegaso^6A-6Dα⁻^), and Wasko^ω⁻^, revealed significant differences (P < 0.001) in the abundance of individual gliadin proteins (α-, ω-, and γ-gliadins) as well as in the combined densitometric estimate of all three gliadin classes (mean value). Pairwise comparisons using Tukey’s HSD post hoc test showed that the mean combined gliadin values differed significantly among all genotypes. However, when the individual gliadin classes were compared among the Pegaso NILs, no significant differences were observed for α-gliadins. Similarly, for the ω- and γ-gliadin fractions, Pegaso^− 1A^ and Pegaso^−1D^ did not differ significantly from each other. In contrast, all other genotype comparisons showed significant differences (Fig. S1).

**Fig. 1.**
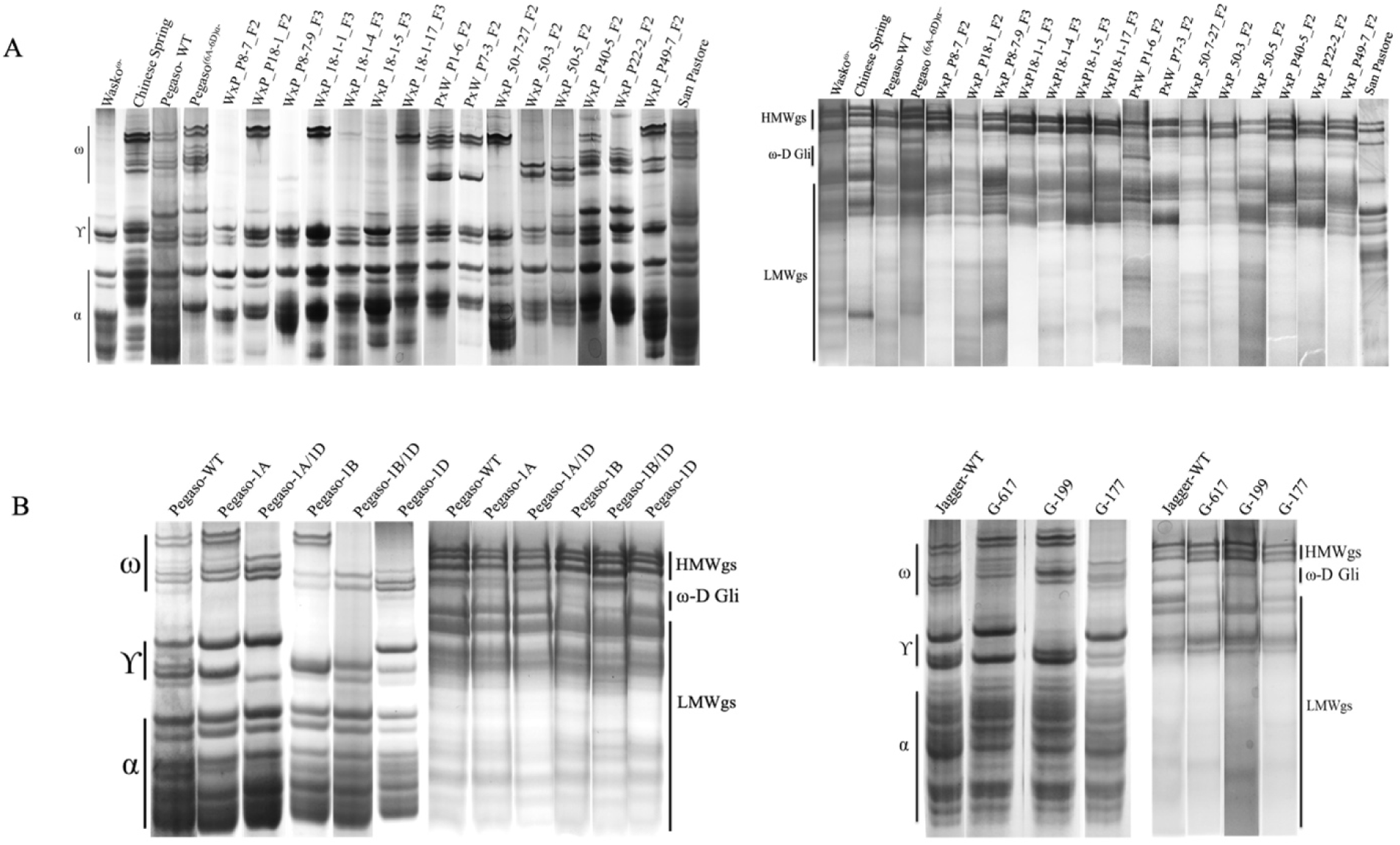
Gliadin and glutenin profiles of different wheat genotypes resolved using acid-urea and SDS-polyacrylamide gels. (A) Gliadin and glutenin profiling of parental genotypes (Wasko^ω-^, Pegaso ^(6A-6D)α-^ , their selected F2 lines, the landrace San Pastor and reference cultivar Chinese Spring (CS). (B) Gliadin and glutenin profiles of Pegaso NILs carrying deletions in one or two *Gli-1* homeoloci and radiation-induced mutants in wheat cv. Jagger background.

Screening of 530 F₂ lines revealed a segregation pattern consistent with expectations for an F₂ population (Fig. S2 and S3; Table S4A and 4B). Detailed examination of banding patterns in the F₂ progenies revealed several noteworthy observations. In two F₂ lines (WxP_P8-7_F2 and WxP_P18-1_F2), a substantial reduction in both α- and ω-gliadins was observed due to successful stacking of mutations from both parents. Among these, WxP-P8-7_F2 lacked the full complement of ω-gliadins and showed a substantial reduction in α-gliadins, retaining only chromosome 6B-specific α-gliadin bands along with one band corresponding to 6D. Another line, WxP_P18-1_F2, carried only two of the seven ω-gliadin bands and lacked most α-gliadins, again retaining only chromosome 6B-specific α-gliadin bands and one 6D specific band (Fig. 1A).

A total of 237F₂ lines were identified that carried deletions in one, two, or three Gli-1 genes (in the homozygous as well as heterozygous state, resulting in the absence of corresponding ω-gliadin bands) in different combinations (182 single deletions, 25 double deletions, and 30 triple deletions). Similarly at *Gli-2* region, 30 lines showed deletion in the D sub genome together with deletion in at least one (out of 3) sub genome whereas 8 lines showed simultaneous deletion in the D sub genome and A sub genome loci (Table S4A 4B). Additionally, three lines carried mutations in four gliadin genes, including both *Gli-1* and *Gli2* loci (quadruple mutants in the homozygous state). Furthermore, when the F₂ line WxP_P8-7_F2 was advanced to the F₃ generation (WxP_P8-7-9_F3), it exhibited a further reduction in α-gliadin content compared to the F₂ generation and closely resembled the α-gliadin profile of the parental line Pegaso^6A-6Dα⁻^. This suggests that the original F₂ line was likely a pentapole mutant with one *Gli-2* locus mutation in the heterozygous state, which segregated in the F₃ generation, enabling the identification of a line combining ω-gliadin deficiency with reduced α-gliadin content.

(ii) *γ-radiation induced mutants in ‘Jagger’ background*: Alignment of whole-genome sequences derived from γ-ray–induced mutants with the Jagger-WT reference enabled the identification of lines carrying deletions in different gliadin genes. These mutants included G-199, which harbors a homozygous deletion on chromosome1B at (0-81.5 Mb), resulting in the deletion of 1 γ-gliadin band and G-617, which contains deletions spanning genes encoding both ω- and γ-gliadins specific to chromosome 1A at (2-6.5 Mb region) in the homozygous state. In another mutant, G-177, a homozygous deletion at chromosome 1D (0–193 Mb) resulted in the loss of ω-gliadin bands 1 and 2, which map to the D subgenome. In addition, this mutant carried a duplication on chromosome 1B (0–407 Mb) and showed reduced γ-gliadin bands when compared to Jagger-WT (Fig. 1B).

(iii) *Group 1, Group 2, and Group 6 nullisomic tetrasomic lines*: The NT lines belonging to group 1 and group 6 chromosomes and N2DT2A exhibited protein deficiencies, while rest of the NT lines for the showed patterns similar to CS. Specifically, Group 1 and Group 6 NT lines enabled the assignment of different protein bands to respective chromosomes. Closer examination of the gliadin banding patterns allowed the assignment of 2 ω-gliadin bands to chromosome 1D, 4 bands to chromosome 1B, and 2 bands to chromosome 1A. Similarly, for γ-gliadins, one band each was assigned to chromosomes 1A, 1B, and 1D. For α-gliadins, 3 bands were assigned to chromosome 6B, 2 bands to chromosome 6D, and 3 bands to chromosome 6A. Remaining two bands (out of 10) correspond to all three group 6 chromosomes (6A, 6B, and 6D) (Fig. S4). For Group 2 NT lines, specifically N2D/T2A (nullisomic for 2D and tetrasomic for 2A), revealed reduction in α-gliadins. Precisely, one out of the 10 α-gliadin bands were found to be missing as indicated in (Fig. S5).

(iv) *Landrace ‘San Pastore’*: When the gliadin banding pattern of this landrace was compared to CS and Pegaso-WT, the ω-gliadin bands showed reduced intensity and one γ-gliadin band specific to chromosome 1D was completely absent (Fig. 1A).

### 3.3. Profiling of high- and low-molecular weight glutenins in different wheat genotypes

There was no change in the profile of HMW glutenin subunits in any of the wheat genotypes analyzed in the present study, as determined by SDS-PAGE and GlutEnSeq. However, LMW glutenin subunits showed reduced accumulation in specific lines, as evidenced by differences in band intensity among the lines. This reduction was observed in selected F_2_ lines, except for W × P_P50-5_F2, P × WxP_P1-6_F2, and W ×P_P18-1_F2. Similarly, among the selected F_3_ lines, all exhibited substantial reductions in LMW glutenin subunit content, except WxP_P8-7-9_F3, which showed only a limited reduction. Similarly, some reduction in LMW-GS was also observed in Pegaso^−1A/1D^. Meanwhile, the landrace ‘San Pastore’ also showed a minor reduction in LMW glutenin subunit content (Fig. 1A-B).

### 3.4. Molecular characterization of Gli1, Gli2, Glu1, and Glu3 loci in selected wheat genotypes using GlutEnSeq

GlutEnSeq analysis of 12 selected wheat lines (Table S1) supplemented the findings from gliadin and glutenin profiling using acid-urea PAGE and SDS-PAGE. To simplify interpretation, deletions at different gliadin (*Gli-A1*, *Gli-B1*, *Gli-D1*, *Gli-A2*, and *Gli-D2*) and glutenin loci (*Glu-A3*, *Glu-B3*, and *Glu-D3*) were summarized under separate headings (see below). Although reduced coverage was observed in many lines, however, only lines carrying deletions of several gliadin genes are discussed in detail. All the heatmaps and/or coverage plots depicting these deletions (based on differences in coverage values) are presented in Figs. 2–4 and Figs. S6–S10.

**Fig. 2.**
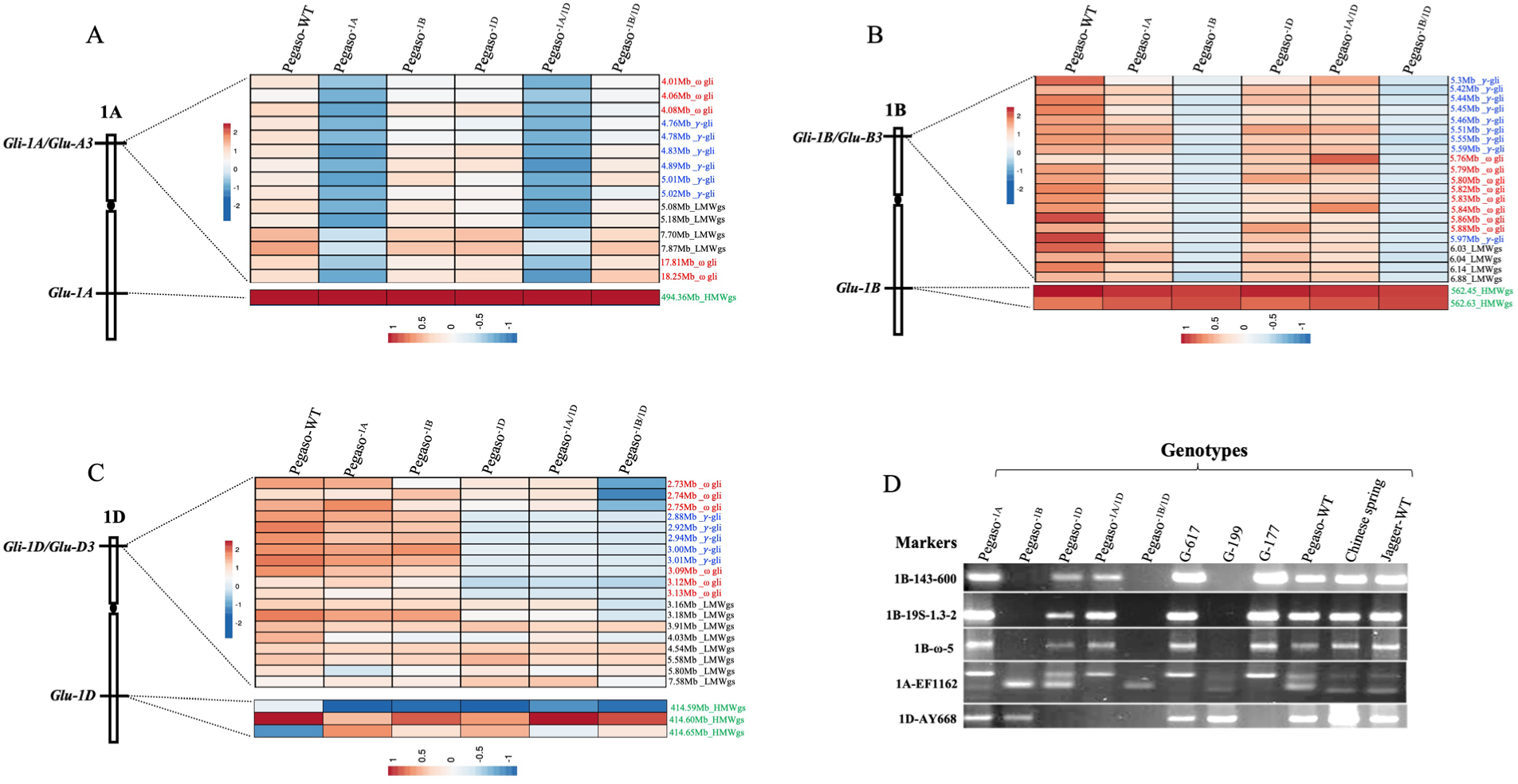
Heatmaps depicting the coverage densities of gluten genes in Pegaso-WT and five Pegaso near-isogenic lines (NILs) across the (A) *Gli-1A/Glu-A3* and *Glu-1A*, (B) *Gli-1B/Glu-B3* and *Glu-1B*, and (C) *Gli-1D/Glu-D3* and *Glu-1D* loci. Gene labels are color-coded according to protein class: ω-gliadins (red), γ-gliadins (blue), low-molecular-weight glutenin subunits (LMW-GSs; black), and high-molecular-weight glutenin subunits (HMW-GSs; green). Rows indicate the physical positions of genes in the Chinese Spring reference genome, whereas columns indicate the genotypes. (D) PCR-based marker analysis of the Pegaso NILs, together with Jagger-WT, Chinese Spring, and three γ-irradiated Jagger lines, using three markers targeting ω-gliadin genes on chromosomes 1A, 1B, and 1D, respectively, as well as two repeat-junction markers designed from the *Gli-1B/Glu-B3* prolamin region specific to chromosome 1B.

**Fig. 3.**
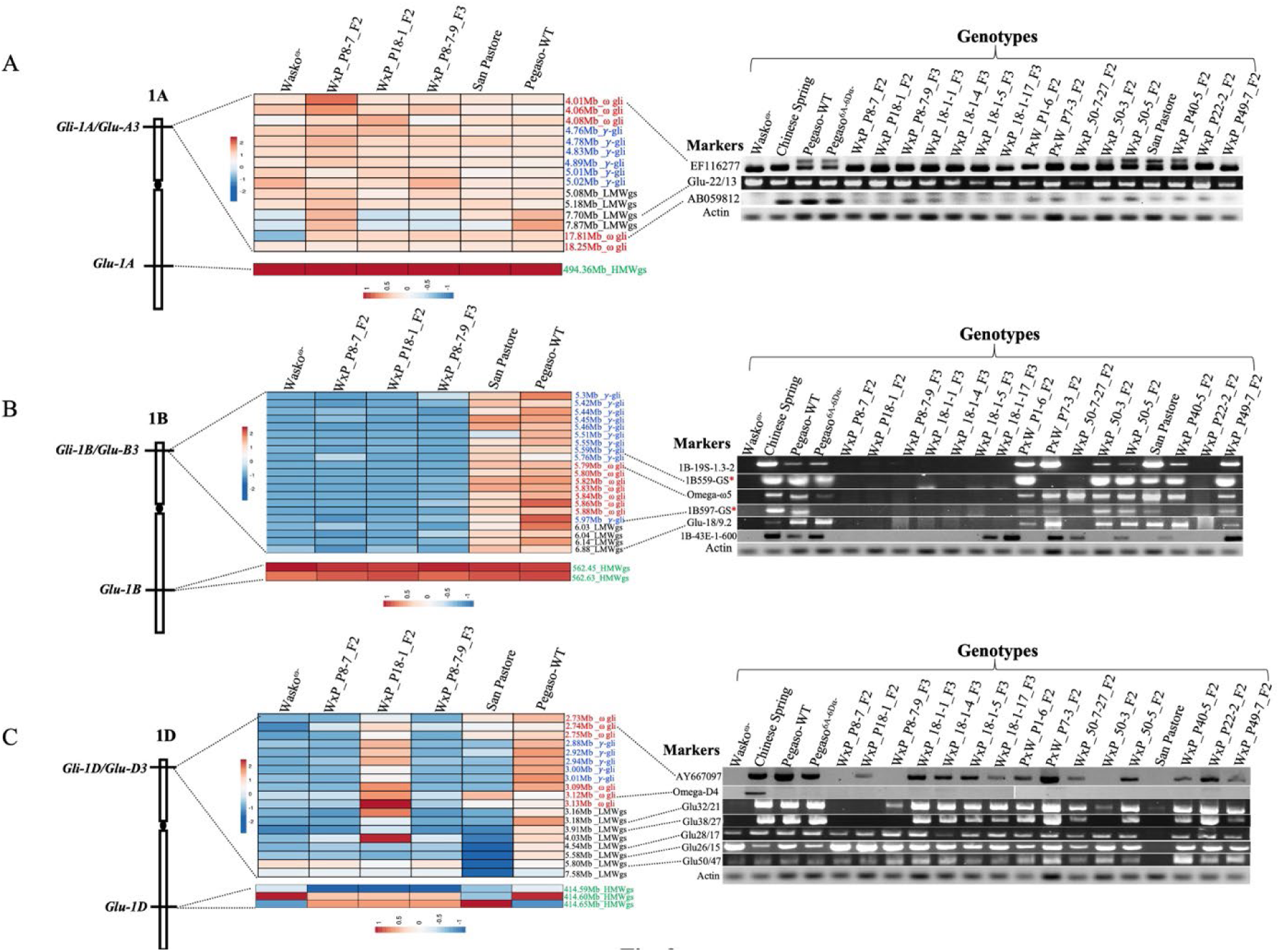
Heatmaps depicting the coverage densities of gluten genes in Pegaso-WT, Wasko^ω⁻^, San Pastore, and three selected progenies derived from the cross Wasko^ω⁻^ × Pegaso^(6A–6D)α⁻^, together with PCR-based marker analysis of the parental genotypes and selected F₂ and F₃ progenies across the (A) *Gli-1A/Glu-A3* and *Glu-1A*, (B) *Gli-1B/Glu-B3* and *Glu-1B*, and (C) *Gli-1D/Glu-D3* and *Glu-1D* loci. Gene labels are color-coded according to protein class: ω-gliadins (red), γ-gliadins (blue), low-molecular-weight glutenin subunits (LMW-GSs; black), and high-molecular-weight glutenin subunits (HMW-GSs; green). Rows indicate the physical positions of genes in the Chinese Spring reference genome, whereas columns indicate the genotypes. Markers denoted by an asterisk (*) were developed in the present study, whereas the remaining markers were previously reported and are listed in Table S2.

**Fig. 4.**
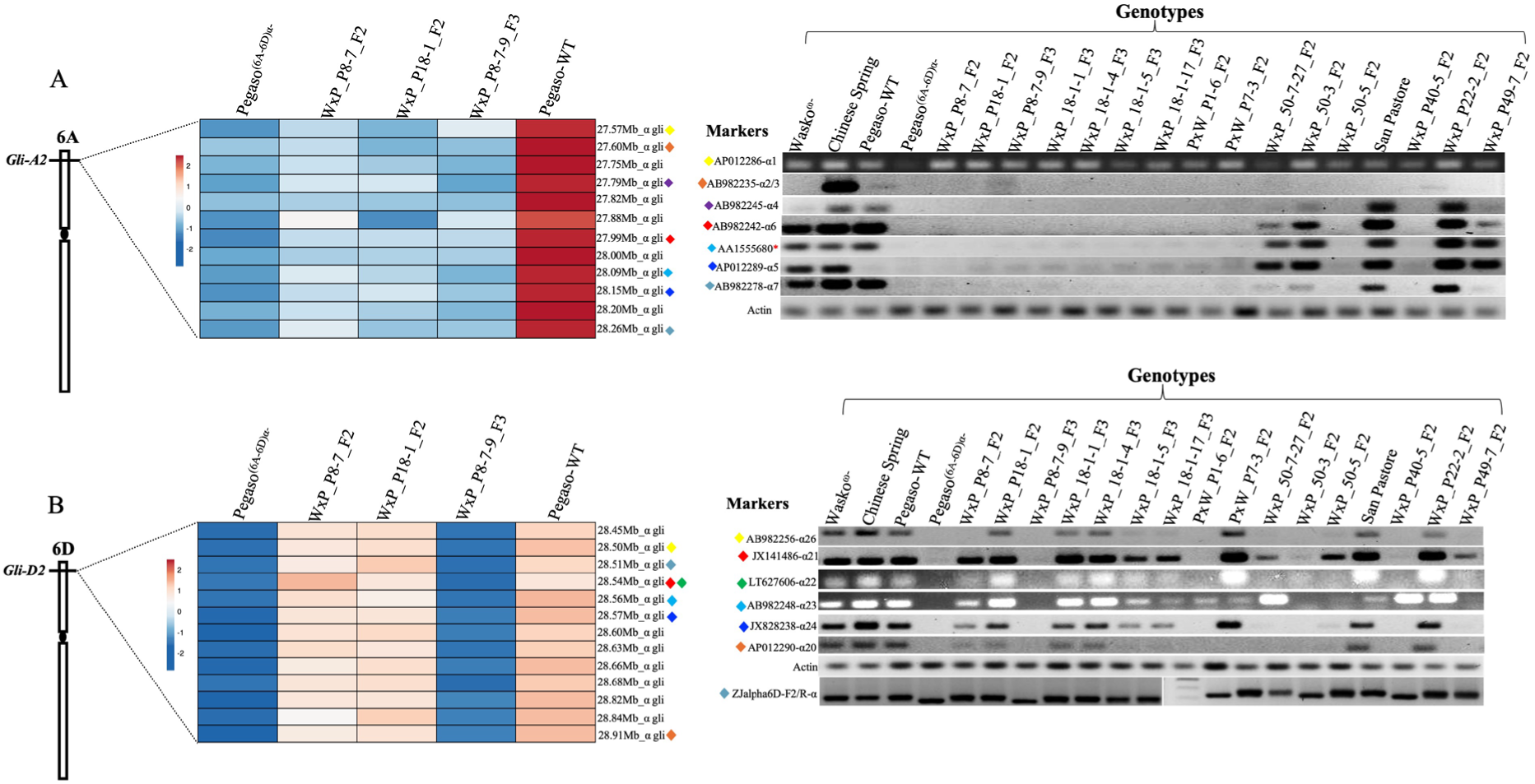
Heatmaps depicting the coverage densities of gliadin genes in Pegaso-WT, Pegaso^(6A–6D)α⁻^, and three selected progenies derived from the cross Wasko^ω⁻^ × Pegaso^(6A–6D)α⁻^, together with PCR-based marker analysis of the parental genotypes and selected F₂ and F₃ progenies across the (A) *Gli-A2* and (B) *Gli-D2* loci. Rows indicate the physical positions of genes in the Chinese Spring reference genome, whereas columns indicate the genotypes. Diamond symbols link each marker shown on the agarose gels to its corresponding physical position within the *Gli-A2* or *Gli-D2* locus. Markers denoted by an asterisk (*) were developed in the present study, whereas the marker Zjalpha 6D-F2/R ZJ targets a 33-mer epitope. The remaining markers were previously reported and are listed in Table S2.

*Gli-A1/Glu-A3* (*4-8Mb and 17-19Mb*)*, and Glu-A1* (*494.3Mb-494.8Mb*): A total of 15 genes map to the *Gli-A1/Glu-A3* locus, including 5 ω-gliadin, 6 γ-gliadin, and 4 LMW-GS genes. The *Glu-A1* locus in Pegaso-WT carried a single HMW gene (1Ax null) and therefore exhibited no change in coverage (Figs. 2A, 3A and S7A). A complete deletion of 12 genes out of 15 gluten genes at the *Gli-A1/Glu-A3* locus—including 6 γ-gliadin genes, 2 ω-gliadin genes, and 4 LMW-GS genes—was observed in Pegaso^−1A^ and Pegaso^−1A/1D^ lines (Fig. 2A and S6A). A few interstitial deletions (or regions of very low coverage) involving LMW-GS genes were also observed at 7.70–7.87 Mb in Wasko^ω−^ and the F_2_ line WxP_P18-1_F2, and at 7.19–7.87 Mb in San Pastore. Additionally, Wasko^ω−^ showed deletion of one of the five ω-gliadin genes at 17.8 Mb region, as depicted in the coverage plot (Fig. 3A and Fig. S7B).

*Gli-B1/Glu-B3* (*5Mb-7Mb*) *and Glu-B1* (*562.4-562.6Mb*): The *Gli-B1*/*Glu-B3* region consists of 8 ω-gliadin genes, 9 γ-gliadin genes, and 4 LMW-GS genes, while the *Glu-B1* locus contains two HMW-GS genes with two subunits (1Bx7 and 1By9). Similar to chromosome 1A, no deletion spanning the *Glu-B1* locus on chromosome 1B was observed (Figs. 2B, 3B and S8A). All genes within the *Gli-B1* locus showed substantially low coverage in Pegaso^−1B^ and Pegaso^−1B/1D^ lines, indicating complete deletion of all genes at this locus in these lines (Figs. 2B and S6B). As expected, Wasko^ω−^ and the three progeny plants from the Wasko^ω−^ × Pegaso^(6A–6D)α−^ cross (WxP_P8-7_F2, WxP_P18-1_F2, and WxP_P8-7-9_F3) showed complete elimination of these genes (Fig. 3B and Fig. S8B).

*Gli-D1/Glu-D3* (*2-7.6 Mb*) *and Glu-D1* (*414.5 Mb-414.7Mb*): *Gli-D1* encompasses a total of 19 gluten genes, including 6 ω-gliadin genes, 5 γ-gliadin genes, and 8 LMW-GS genes. Much like chromosomes 1A and 1B, the *Glu-D1* locus on chromosome 1D encompasses three HMW-GS genes (1Dx5 and 1Dy10) and showed no loss of genes observed in any of the studied genotypes (Fig. 2C, 3C and S9A). Analysis of sequences across 12 selected genotypes revealed deletions spanning most genes in Pegaso^−1B/1D^ (from 2.73 Mb to 3.18 Mb, covering 6 ω-gliadin genes, 5 γ-gliadin genes, and 2 LMW-GS genes; Figs. 2C and S6C), Wasko^ω-^, and the two progeny plants, WxP-P8-7_F2 and WxP-P8-7-9_F3 (Fig. 3C and Fig. S9B). Although the lack of coverage observed for a few genes was more pronounced in Wasko^ω-^ compared to Pegaso^−1B/1D^. Sequence analysis of San Pastore showed a complete deletion of the region containing all five γ-gliadin genes and two ω-gliadin genes. Interestingly, the landrace ‘San Pastore’also showed substantially reduced coverage across all LMW-GS genes, with complete deletion of at least four LMW-GS genes (Fig. 3C and Fig. S9B). This represents the first characterization of a *Gli-D1/Glu-D3*-deficient biotype in San Pastore, as previous studies have only reported the characterization or isolation of *Gli-B1/Glu-B3*-deficient biotypes, which we have also recovered.

*Gli-A2* (*Ch 6A: 27.55-28.22Mb*) *and Gli-D2* (*6D: 28.4Mb-28.9Mb*): The *Gli-A2* and *Gli-D2* regions span 12 and 13 α-gliadin genes, respectively. Since these genes are located on group 6 chromosomes, only Pegaso^(6A–6D)α−^, Pegaso-WT, and the selected progeny from the Wasko^ω−^ × Pegaso^(6A–6D)α−^ cross were examined for coverage across this gene space. As expected, the entire region containing 25 α-gliadin genes showed low coverage profiles in Pegaso^(6A–6D)α−^, with 22 genes showing complete deletion. Interestingly, the F_3_ progeny (WxP-P8-7-9_F3) largely resembled the deletion pattern observed in Pegaso^(6A–6D)α−^, whereas the two F_2_ lines (WxP_P8-7_F2 and WxP_P18-1_F2) showed only substantially reduced coverage in the chromosome 6A region encompassing the 12 α-gliadin genes. This likely reflects the heterozygous status of the chromosome 6D deletion in the F_2_ lines. As expected, Pegaso-WT did not show any reduction in coverage in these regions (Figs. 4A, 4B, and Fig. S10A-B).

### 3.5. Validation of GlutEnSeq results using molecular markers

The deletions identified using GlutEnSeq were validated using 30 different PCR-based markers. In addition to the lines used for GlutEnSeq, a few F_2_ and F_3_ lines were also included in the marker analysis. Of the 30 markers, 3 were specific to *Gli-A1/Glu-A3*, 6 to *Gli-B1*, 7 to *Gli-D1*, 7 to *Gli-A2*, and 6 to *Gli-D2*, while one marker was specific to the 33-mer epitope region located at *Gli-D2* (Figures 2–4; see Table S2 for marker details). These markers confirmed the deletions identified through gliadin profiling and GlutEnSeq in all studied lines, including Pegaso^(6A-6D)α-^, Wasko^ω-^, and the derived F_2_ and F_3_ lines. Among the 30 markers, four—1B559 and 1B597 (specific to *Gli-B1*, γ-gliadins), AA1555680 (*Gli-A2*), and ZJalpha6D-F2/R (*Gli-D2*)—were developed in this study. Except for ZJalpha6D-F2/R, which showed a size difference, all other markers detected the presence or absence of amplification products in cases of deletion. Additionally, because the ZJalpha6D-F2/R marker targets the 33-mer epitope, it clearly differentiated Pegaso^(6A-6D)α-^, two F_2_ lines, and one F_3_ line from the remaining genotypes. Interestingly, two primer sets—1B-43E-1-600-Repeat Junction and 1B-19S-1.3-2-Repeat Junction—designed from the prolamin region of *Gli-1B* and *Glu-B3* revealed an approximately 5.8 Mb deletion (out of a 6.5 Mb region) encompassing LMW-GS, omega-5 gliadin, and gamma gliadin genes in Wasko^ω-^ and 9 F_2_ lines. Additionally, these markers also confirmed deletions of specific *Gli1/Glu3* loci detected through gliadin profiling and GlutEnSeq in specific sub-genomes of the Pegaso NILs and gamma ray-induced Jagger mutants (Fig. 2D). Specifically, markers for *Gli-1* loci were used because these lines carry deletions at these loci. As expected, the absence of amplicons in these lines validated the deletions identified through GlutEnSeq and/or gliadin profiling. Briefly, markers specific to the *Gli-1B* locus—Gli-1B-143-600F/R, Gli-1B19S1.3F/R, and Gli-1Bω-5F/R—showed no amplification in Pegaso^−1B^ and Pegaso^−1B/1D^, validating our initial analysis using GlutEnSeq and gliadin profiling, as well as in one gamma-radiated mutant (G-199), confirming the gliadin profiling results (Fig. 2D). Similarly, the *Gli-1A*-specific marker Gli-1AEF1162F/R showed no amplification in Pegaso^−1A/1D^, G-617, and G-177, while the *Gli-1D*-specific marker AY667 F/R showed no amplification in Pegaso^−1D^, Pegaso^−1A/1D^, Pegaso^−1B/1D^, and G-177.

### 3.6. Gliadin gene-product relationship based on molecular and biochemical data

In the present study, for deducing the relationship between protein bands and the corresponding genes, we prepared a reference protein map based on all our gliadin profiles (Figs. 1A and 1B; Fig. S4) and taking support from earlier literature and accordingly, the protein bands were assigned to the wheat sub-genomes A, B and D (Fig. 5A). Using this reference, the present study revealed the following relationships between the gliadin genes and the corresponding protein bands: (i) ω-gliadins: A total of eight protein bands were resolved using acid-urea PAGE. The gene-to-protein relationship showed that two protein bands assigned to chromosome 1A corresponded to a single gene (suggesting protein isoforms), four protein bands assigned to chromosome 1B corresponded to eight genes (suggesting lack of resolution or presence of pseudogenes), and the remaining two protein bands assigned to chromosome 1D corresponded to four genes (also suggesting lack of resolution or presence of pseudogenes). (ii) γ-gliadins: A total of four protein bands were resolved using acid-urea PAGE. The gene-to-protein relationship revealed that two protein bands corresponded to six genes on chromosome 1AS, one protein band corresponded to eight genes on chromosome 1BS, and one protein band corresponded to five genes on chromosome 1DS (suggesting lack of resolution or presence of pseudogenes). (iii) LMW-GS: Correspondence based on protein bands could not be established; however, our GlutEnSeq analysis provided evidence for both genes and pseudogenes. Specifically, four LMW-GS genes were assigned to chromosome 1A and 2 out of these 4 genes showed reduced coverage in Wasko^ω-^, Pegaso^−1A^, and Pegaso ^−1A/1D^. On 1BS, four LMW-GS genes were assigned, and all of them showed reduced coverage in all the mutants except ‘San Pastore.’ On 1DS, eight LMW-GS genes were assigned; however, the coverage differed for all the eight genes in different mutants analyzed. For instance, all the eight LMW-GS showed reduced coverage in San-Pastore, while only 5 (out of 8) showed reduced coverage in Pegaso^−1B/1D^ and 2 (out of 8) genes showed reduced coverage in Pegaso^−1D^ and Pegaso^−1A/1D^. All these results were based on either complete deletion or reduced coverage observed in the GlutEnSeq coverage profiles and molecular marker analysis (Fig. 5B; Figures 2–4; Figs. S6–S10). However, LMW-GS bands could not be completely resolved on Bio-Rad Mini-PROTEAN SDS-PAGE gels (Fig. 1A and Fig. 1B), and therefore, further analysis is needed to establish the complete LMW-GS gene-protein relationship. (iv) α-gliadins: A total of 10 protein bands were resolved using acid-urea PAGE. The gene-to-protein relationship indicated that three protein bands corresponded to 12 genes on chromosome 6AS, while two protein bands corresponded to 13 genes on chromosome 6DS, suggesting lack of resolution or presence of pseudogenes. Three protein bands were mapped to chromosome 6BS. The remaining two (out of 10) protein bands may correspond to genes from all three sub-genomes, as these protein bands appeared to map to all three sub-genomes (Fig. 5C).

**Fig. 5.**
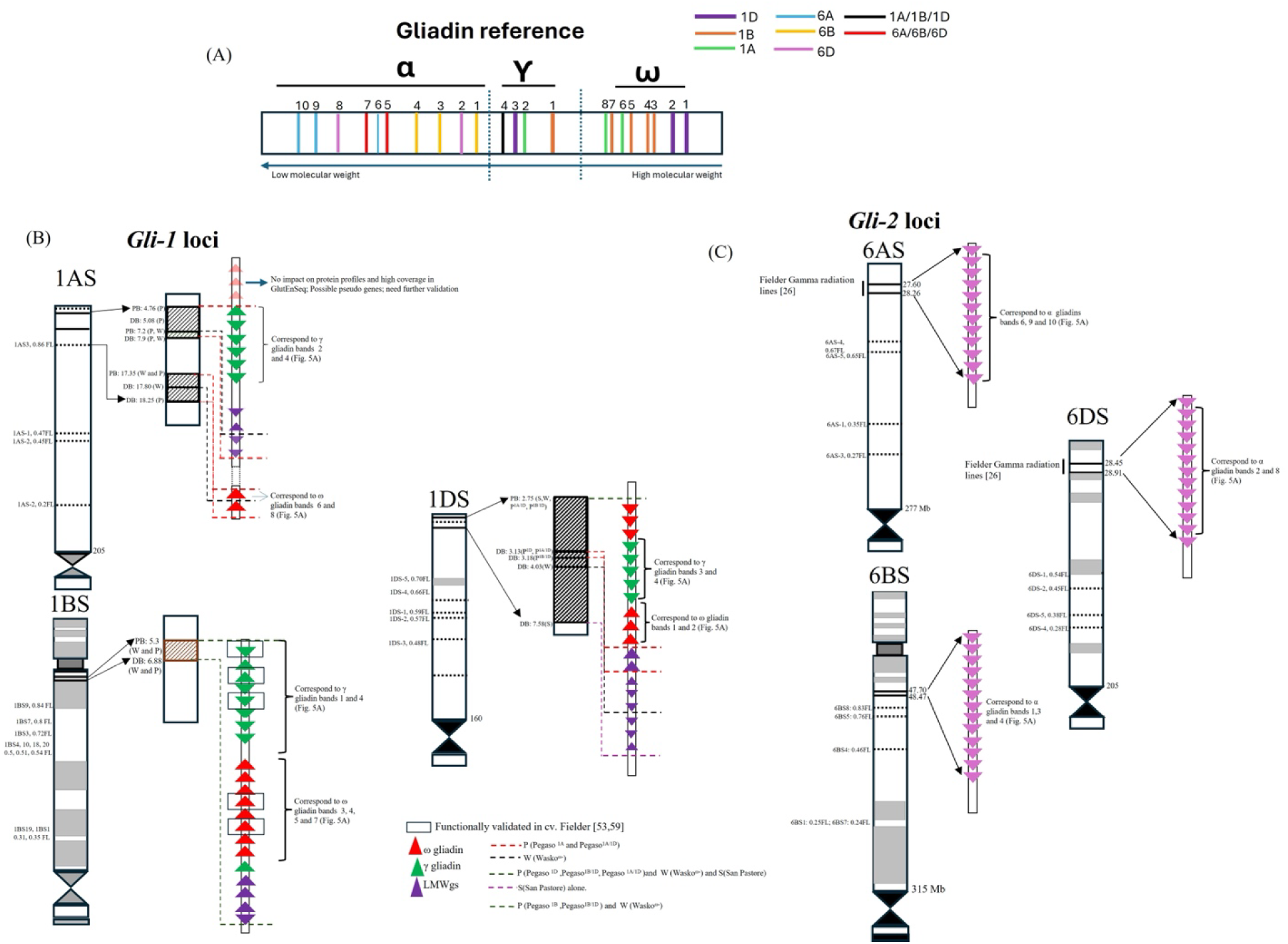
Physical mapping and gene–protein relationships of the *Gli1* and *Gli2* loci were established based on information from the available literature [54–55,32] and genetic stock screening performed in this study using biochemical (acid-urea PAGE and SDS-PAGE), immunological (ELISA), and genomic (GlutEnSeq and DNA marker) methods. (A) A reference gliadin map showing the number and distribution of ω-, γ-, and α-gliadin protein bands resolved by acid-urea PAGE for the different genotypes studied. The thickness of each bar represents the intensity of the corresponding protein band observed in the acid-urea PAGE profile. (B) Physical mapping of the *Gli-1* loci and establishment of ω-gliadin, γ-gliadin, and LMW-GS gene-to-protein relationships based on GlutEnSeq analysis. (C) Physical mapping of the *Gli-2* loci and establishment of α-gliadin gene-to-protein relationships based on GlutEnSeq analysis. The correspondence between genes and protein bands was determined using GlutEnSeq data (Figs. 2, 3 and 4; Figs. S6-10) and acid-urea PAGE profiles (Figs. 1A, 1B and Fig. S4). Breakpoints carried by interstitial and terminal deletion mutants are indicated by shaded blocks along each chromosome. These breakpoints were identified through our GlutEnSeq analysis and supported by previous studies [32,54]. Fraction length (FL) values are based on CS deletion lines [55]. PB and DB denote proximal and distal breakpoints, respectively. In Fig. 5B, the letters W, P, and S in parentheses following each PB and DB indicate breakpoints identified using Wasko^ω−^, one or more Pegaso near-isogenic lines (Pegaso^−1A^, Pegaso^−1B^, Pegaso^−1D^, Pegaso^−1A/1B^, and Pegaso^−1B/1D^), and the landrace ‘San Pastore,’ respectively. In Figure 5C, information generated using Pegaso^(6A–6D)α−^ was used as the reference for gene mapping.

### 3.7. Immunological analysis of selected wheat lines

Gliadin levels estimated by ELISA (in parts per million, ppm) were expressed as a percentage relative to the CS estimate. In the present study, the gliadin content of CS was determined to be 19,044 ppm (treated as 100%). Accordingly, the gliadin content of the two parental lines, Wasko^ω−^ and Pegaso^(6A–6D)α−^, was reduced to 33% and 63% of that in CS, respectively. Similarly, among the eight F_2_ lines evaluated by ELISA, four lines showed a reduction in gliadin content ranging from 64% to 50%, whereas the remaining four lines exhibited reductions between 37% and 11%. Among the three F_3_ lines, two showed reduced gliadin content of 59% and 56%, while the third line showed a reduction to 38%. Overall, five progenies exhibited gliadin content reduced to less than 50%, as shown in Fig. 6A.

**Fig. 6.**
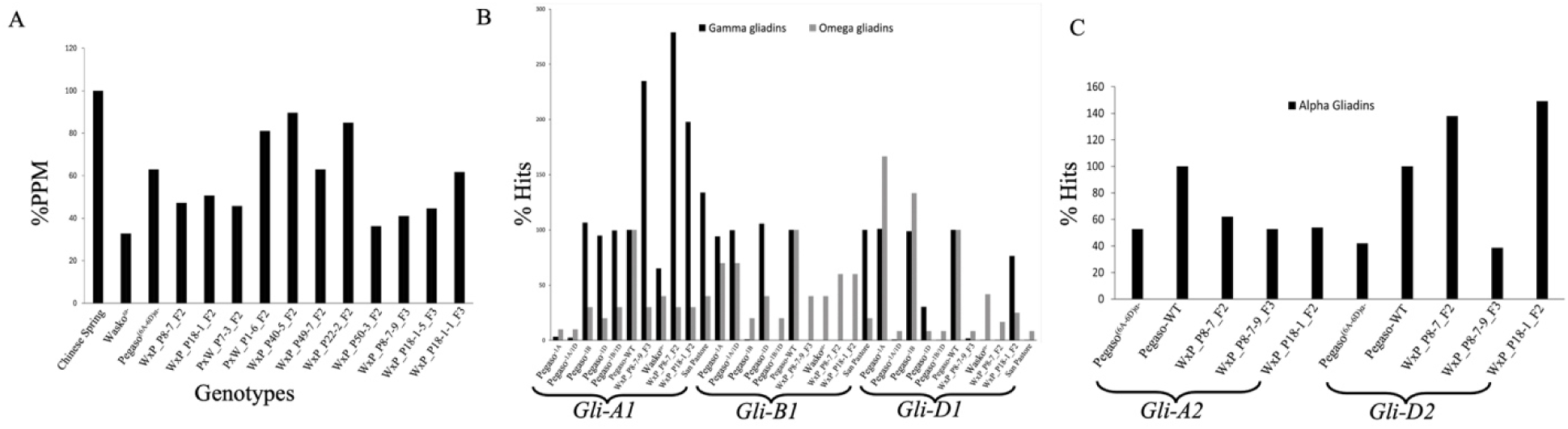
Immunological evaluation of the reduced-gluten wheat lines. (A) Results of ELISA using the R5 antibody, (B) *in silico* epitope load (%) analysis using GlutEnSeq reads at the *Gli-A1*, *Gli-B1*, and *Gli-D1* loci, (C) *in silico* epitope load (%) analysis using GlutEnSeq reads at the *Gli-A2* and *Gli-D2* loci.

### 3.8. In silico analysis of immunogenic epitopes in selected wheat lines

The results of the *in silico* epitope analysis revealed notable differences among the five gliadin loci (*Gli-A1*, *Gli-B1*, *Gli-D1*, *Gli-A2*, and *Gli-D2*) examined in this study. For this analysis, the number of hits obtained for Pegaso-WT was considered as 100%, and the hits for other genotypes were expressed as a percentage relative to Pegaso-WT (Figs. 6B and 6C). Using this approach, two genotypes—Pegaso^1A^ and Pegaso^−1A/1D^,—showed a significantly reduced number of hits for *Gli-A1* loci (3%, and 2%, respectively), primarily against the γ-gliadins epitopes DQ2.5 γ1 (PQQSFPEQQ) and DQ8 γ1a (EQPQQPFPQ), whereas the remaining genotypes showed more than 100% hits, except the Waskoω− showed 65% hits for these epitopes. In contrast, for omega gliadin genes at the *Gli-A1* locus, all the 10 lines (excluding Pegaso-WT ) showed fewer than 50% hits, particularly against the epitopes DQ2.5 ω1 (PFPQPEQPF) and DQ2.5 ω2 (PQPEQPFPW), as shown in (Fig. 6B).

For the γ-gliadins at the *Gli-B1* locus, the genotypes Pegaso^1B^, Pegaso^−1B/1D^, and Wasko^ω−^ showed less than 2% epitope hits compared to Pegaso-WT (100%). Similarly, the derived progenies from the cross Wasko^ω−^× Pegaso^(6A–6D)α−^ exhibited the same pattern. However, the landrace ‘San Pastore’ showed 100% hits, indicating either no change or a higher epitope count (Fig. 6B). For the ω-gliadins, reduced epitope hits were also observed in Pegaso^−1B^and Pegaso^−1B/1D^, , with approximately 80% fewer hits compared to Pegaso-WT. Furthermore, Wasko^ω^and one F_3_ line showed a 60% reduction in hits, whereas the Pegaso^1A^ and Pegaso^−1A/1D^ showed more than 70% hits.

The epitope analysis results for the *Gli-D1* locus largely matched the expected patterns; genotypes that were null for this locus showed reduced epitope hits compared to Pegaso-WT. However, among the two F₂ progenies, only one (W × P_P8-7-9_F2) showed a significant reduction in the number ω and γ-gliadins hits, whereas the other (W × P_P18-1_F2) showed a smaller reduction in comparison. Interestingly, ‘San Pastore’ showed the lowest number of hits for gamma gliadin epitopes, confirming the identification of a novel biotype with a significant reduction in gamma gliadins (Fig. 6B).

When compared to the *Gli-A1*, *Gli-B1*, and *Gli-D1* loci, the reduction in epitope hits was less pronounced for the *Gli-A2* and *Gli-D2* loci. Specifically, for *Gli-A2*, epitope hits were reduced to 50–75% in Pegaso^(6A-6D)α⁻^ and the F₃ progeny compared to Pegaso-WT. Similarly, for *Gli-D2*, Pegaso^(6A-6D)α⁻^ and the F₃ line (W × P_P8-7-9_F3) showed reduced hits (42% and 37%, respectively), whereas the remaining genotypes showed higher numbers of hits compared to Pegaso-WT. This pattern corresponded with the gliadin profile, indicating the loss of key DQ2.5 and DQ8 alpha gliadin epitopes. Interestingly, the key 17-mer peptide was also absent in the parental genotype Pegaso^(6A-6D)α⁻^ as well as in the F₃ progeny (Fig. 6C).

### 3.9. Seed attributes of reduced-gluten wheat lines and wild type

The 100-seed weight of Pegaso-WT, its mutants, and Wasko^ω−^ ranged from 3.16 g (Pegaso^(6A-6D)α−^) to 3.86 g (Wasko^ω−^), indicating non-significant variation. Similarly, different seed size parameters, including seed area, seed length, seed width, length-to-width ratio, and perimeter, were recorded. As expected, the highest seed area was observed in Pegaso-WT (1285.69 mm²), whereas the lowest was recorded in W × P_P8-7-9-F3 (754.13 mm²). The seed length-to-width ratio did not show much variation and ranged from 2.25 (Pegaso^−1A/1D^) to 2.84 (W × P_P8-7-9-F3). Overall, all seed size parameters were lowest in the progeny (W × P_P8-7-9-F3) compared to the parental genotypes (Fig. S11; Table S5).

## 4. Discussion

Wheat serves as a staple food for approximately 36% of the global population, and projections indicate that worldwide consumption may exceed 800 MMT in the near future. The continued rise in wheat consumption, particularly in regions that have not traditionally been major consumers—highlights an increasing vulnerability of populations to wheat-related disorders. This trend underscores the urgent need to develop wheat lines with reduced levels of immunogenic proteins to improve food safety by lowering the risk of accidental exposure–triggered immunological reactions. Considerable efforts have been made toward this goal; however, the most promising lines developed to date have largely relied on RNA interference or gene-editing approaches [41–43,5]. Despite their effectiveness, such materials, especially those derived from genetic transformation—face significant regulatory and public acceptance challenges. Recent survey data further indicate that even when gene-edited crops are approved, their acceptance varies depending on the crop and target population, with reduced-gluten wheat receiving relatively limited support [23].

In contrast, materials developed through conventional crossbreeding are generally more widely accepted. However, the majority of previously characterized lines have focused on modifying a single gliadin class, such as α-gliadins [26,25] or ω-gliadins [24]. Efforts targeting multiple gliadin classes have been limited and, in some cases, discontinued [27]. As a result, no currently available lines effectively eliminate major immunogenic gliadins or sufficiently reduce immunogenic epitopes, leaving a critical gap in available germplasm.

In this study, we addressed this gap by first characterizing available germplasm (Table S1) using both protein profiling and genomic analyses. We then performed crosses between two mutant lines, Wasko^ω⁻^ and Pegaso^(6A–6D)α-^, with the objective of stacking desirable mutations affecting immunogenic gliadins into a single genetic background. Although these mutant lines had been previously evaluated using biochemically (PAGE and RP-HPLC; [25,24]and immunologically [24,44–45], they had not been utilized for mutation stacking despite harboring valuable alleles at the *Gli1* and *Gli2* loci. Similarly, Pegaso near-isogenic lines (*Gli-1* single nulls - Pegaso^−1A^, Pegaso^−1B^, and Pegaso^−1D^, and *Gli-1* double nulls Pegaso^−1A/1D^ and Pegaso^− 1B/1D^) have been characterized biochemically for phenolic content [46] and gluten composition using LC-MS [47]. However, none of the mutants used in this study had been characterized at the genomic level. Such characterization is essential for the development of molecular markers to facilitate breeding efforts, including gene pyramiding, transfer of mutations into adapted backgrounds, and efficient screening of germplasm for reduced immunogenicity. Therefore, this study serves two primary purposes: (i) to establish a foundational understanding linking specific gliadin genes to their protein products, and (ii) to develop genetic and molecular resources that enable the breeding of reduced-gluten wheat lines through conventional approaches. Collectively, these efforts contribute to bridging the gap between consumer acceptance and the development of safer wheat varieties.

### 4.1. Identification of novel trans-regulators of wheat gluten protein accumulation

Protein profiling of the group 2 NT line in the CS background, specifically N2D/T2A (nullisomic for 2D and tetrasomic for 2A), revealed a noticeable reduction in α-gliadins, which was unexpected given that known α-gliadin genes are located on group 6 chromosomes (Fig. S4). These results suggest that the expression of group 6 α-gliadin genes may be regulated either through trans-acting factors, such as transcriptional regulators, or through epistatic interactions involving genes on group 2 chromosomes. Similar observations were reported by [48] and [49], who proposed a role for chromosomal imbalance or epistatic regulation. More recently, [50], provided further support for this epistatic or trans-regulatory control of α-gliadins. In particular, chromosome 2A harbors the transcriptional repressor *TaB3-2A1*, which binds to specific cis-regulatory modules, including RY-like elements, represses gluten gene transcription, and shifts metabolic resources toward starch biosynthesis. This mechanism also supports the chromosome imbalance hypothesis proposed by [48], since the extra dosage of 2A in the N2D/T2A line could suppress specific gluten proteins. Further validation of this model could be achieved through transgenic overexpression of *TaB3-2A1*.

### 4.2. Integration of genomic and protein profiling reveals gene–protein relationships in wheat gliadins

One of the objectives of this study was to assign specific gluten genes and protein bands (this study), HPLC peaks, and mass spectra peaks [47], to sub-chromosomal bins and establish their correspondence. Using cytogenetic stocks, including nullisomic tetrasomic lines for specific chromosomes, terminal chromosomal deficiencies (CS deletion lines), and interstitial chromosome deletions (mutants in Wasko^ω-^, Pegaso, and Jagger backgrounds), we were able to map gliadin loci to sub-telomeric regions (bins) on group 1 chromosomes (ω- and γ-gliadins, as well as LMW-GS and HMW-GS) and group 6 chromosomes (α-gliadins). Furthermore, GlutEnSeq reads mapped to the CS reference assembly enabled determination of proximal and distal breakpoints of interstitial chromosomal deletions, allowing us to localize gliadin genes to different sub-telomeric regions (group 1: 4-8Mb and 17-18Mb on 1AS; 5.3-7Mb on 1BS; 2.5-4.5Mb on 1DS; group 6: 27.5-28.5Mb on 6AS , 47.7-48.47Mb on 6BS and 28-29Mb on 6DS). Earlier studies reported broad genomic locations of gliadin genes [51–53], and more recent sequencing efforts refined their localization [34,54,7,26,8]. However, none of these studies established direct correspondence between genes and their protein products. Based on our results (CS and all Pegaso mutants) and previous reports [55],[32], [54], we mapped seven gliadin protein bands (2 ω, 2 γ, and 3 α) to the A subgenome, ten bands (4 ω, 1 γ, and 3 α) to the B subgenome, and four bands (2 ω, 1 γ, and 2 α) to the D subgenome (Fig. 5A; Fig. 1A, 1B and Fig. S4). Acid-urea PAGE sufficiently resolved different gliadin subfamily members, and the numbers were consistent with the 38 protein peaks identified in Pegaso-WT [47]. Among these peaks, six were assigned to the *Gli-A1* locus, six to the *Gli-B1* locus, and five to the *Gli-D1* locus. These values are also consistent with [30]. In certain cases, higher resolution is desirable, which can be achieved using 2D-PAGE. This method has identified 39-69 major protein spots, including 23–49 α-gliadins, 11–13 γ-gliadins, and 5– 7 ω-gliadins in Butte86 [56]. However, 2D-PAGE is low throughput, limiting its application for large numbers of genotypes.

In support of our findings, gliadin genes on chromosome 1BS were also mapped in an earlier study, and their physical positions were consistent with our analysis [32]. In that study, gamma radiation–induced deletion lines of cv. Paragon were subjected to GlutEnSeq analysis, and deletions were correlated with protein bands based on changes observed through acid-PAGE profiling (Fig. 5A and Fig. 5B). Similarly, α-gliadin–encoding genes on chromosomes 6A and 6D were shown to be mapped to homologous regions in wheat cv. Fielder, based on deletions observed in gamma radiation–induced mutants [26]. In our study, protein profiles showed complete deletion of bands corresponding to the loss of all 11 genes on chromosomes 6A and 6D. However, in an earlier study, as mentioned, not all α-gliadins appeared to be deleted [26]. This discrepancy may be attributed to genotypic differences between Fielder and Pegaso mutants or to the heterozygous nature of the double mutant. Furthermore, in another study by [54], CRISPR/Cas9-based gene-edited mutants were generated in cv. Fielder for ω- and γ-gliadins. Functional genes were identified based on the reduction in intensity or deletion of corresponding protein bands. Specifically, three and two ω-gliadin genes on chromosomes 1AS and 1BS, respectively, and four and three γ-gliadin genes on the same chromosomes were identified as functional genes based on the observed protein deletions. However, since the genotypes analyzed in the present study differ from those in previous studies (Table S6), it is difficult to definitively identify functional genes without complete genome sequencing and expression analysis. Nevertheless, the results clearly identified pseudogenes in almost all cases. Therefore, future studies involving whole-genome sequencing and RNA-Seq are planned to elucidate the functional roles of these genes in the studied germplasm.

### 4.3. In silico epitope analysis enabled assessment of immunogenic potential of reduced-gluten wheat lines

The deletion of immunogenic gliadins was also supported by epitope analysis. *In silico* analysis suggested that stacking these deletions markedly reduced immunogenic epitopes, including the key 33-mer and DQ2.5-glia-α1/α2 motifs, which are the two major CD epitopes [57]. Similarly, the loss of ω- and γ-gliadins substantially reduced immunogenic epitopes across the *Gli-1* locus. Earlier studies involving RNAi and CRISPR approaches [54,43,5,41–42,58] also reported significant reductions in one or more gliadin epitopes involving both the *Gli-1* and/or *Gli-2* loci, sometimes extending beyond the observations of the present study. Some of these differences may be explained by variations in the analytical methods used. The GlutEnSeq analysis indicates that the genotypes developed in this study, carrying multiple null mutations, showed a drastic reduction in the overall immunogenic potential of storage proteins. Previous studies have shown that motifs derived from ω-gliadin, such as PFPQPEQPFPW, and the α-gliadin 33-mer are major drivers of T-cell activation in celiac disease [57], and that most of these epitopes are encoded by the D sub-genome chromosomes 1D and 6D [34]. In addition to α- and ω-gliadins, γ-gliadins are also major contributors to celiac disease onset. The primary γ-gliadin epitopes include DQ2.5-glia-γ5, DQ2.5-glia-γ2, DQ2.5-glia-γ4f, and DQ2.5-glia-γ1, which collectively represent a substantial portion of wheat’s immunogenic potential [59–60], and are also significantly reduced in the reduced-gluten lines developed in this study.

A similar study was conducted by [60], using RNA-seq. This study showed that Gli-γ1-1D and Gli-γ2-1B are the two most highly expressed γ-gliadin genes, and their knockdown resulted in an almost 70% reduction in epitope load. Interestingly, the San Pastore biotype analyzed here, as well as the single- and double *Gli1*-null mutants in Pegaso background, also showed a drastic reduction in γ-gliadin epitopes, especially DQ2.5-glia-γ5. Wasko^ω⁻^ was also previously characterized for its immunogenic potential [24], where it showed no detectable IgE binding to ω-gliadin (compared to the control) when tested against sera from wheat-allergic individuals. Similarly, peptic-tryptic digest of Gli-A2 deficient ‘Raeder’ (*Gli-A2* null allele doner of Pegaso^(6A–6D)α−^) and Gli-D1 and Gli-A2 deficient line, C173 was assessed using celiac mucosa in an in vitro organ culture system and these lines exhibited a significantly lower toxicity in respect of bread wheat with a full gliadin complement [44–45]. Moreover, the absence of immunological reactivity due to the loss of epitopes in Wasko^ω⁻^, Gli-A2 deficient Raeder, and C173 validated the findings of the *in silico* analysis. In alignment with these studies, our material also exhibited similar reductions in all three major gliadin epitopes across the *Gli-1* and *Gli-2* loci, confirming that stacking null mutations of ω-, γ-, and α-gliadins can significantly reduce immunogenic potential.

### 4.4 Developing resources for reduced-gluten wheat breeding

The present study resulted in important genomic and genetic resources for breeding wheat with reduced immunogenic potential without significantly compromising end-use performance. The mutants (Wasko^ω⁻^ and Pegaso^(6A–6D)α⁻^) are certainly important materials for reduced gluten content, as revealed through protein profiling and GlutEnSeq. Advanced lines were developed by crossing these mutants. The reduction in gliadin content could be further enhanced by top-crossing these lines with mutants showing reductions in gamma gliadins (in Pegaso and Jagger backgrounds) and a *Gli-B2* mutant line [26]. As expected, the F₂ and F₃ lines generated by crossing Wasko^ω⁻^ and Pegaso^6A-6Dα⁻^ were deficient in major gliadins (group 1, ω- and γ-gliadins and group 6, α-gliadins), including the major celiac disease epitopes, as revealed through bioinformatics analysis. Furthermore, our GlutEnSeq analysis revealed the presence of intact HMW-GS, which are known to be important for maintaining bread-making quality. Additionally, the glutenin-to-gliadin ratio in these lines, relative to the wild type, shifted toward higher glutenin content, which is generally associated with improved end-use performance [47]. However, a search for genes in the regions encompassing the deleted gliadin genes revealed 11–100 additional genes (excluding gliadin genes) encoding a variety of gene classes (Tables S7–S11). To precisely determine the number of non-gliadin genes deleted and to define the deletion boundaries, in-depth sequencing of the mutant lines is required; this work is currently ongoing. We also recognize that these mutants must undergo agronomic characterization to ensure there is no penalty in yield attributes and/or increased susceptibility to environmental or biological stressors. Similarly, although the presence of HMW-GS was confirmed through *in silico* analysis and SDS-PAGE (Figs. 1-3 and Figs. S7-S9), bread-making quality still needs to be experimentally validated. Due to the limited seed availability and the preliminary stage of the experiments, this could not be assessed in the present study. Furthermore, additional mutants (Pegaso^−1A/1D^ and Pegaso^−1B/1D^) are being incorporated into future crossing programs to stack these mutations in the F₂ and F₃ lines developed in this study. A work plan outlining future experiments is presented in (Fig. S12).

Data on seed weight and seed size rarely revealed any drastic variation within the parental mutants and Pegaso-WT. However, the progeny line (W × P_P8-7-9_F3) showed relatively lower values for both 100- seed weight (2.8 g, whereas the remaining lines ranged from 3.16 to 3.86 g) and seed size parameters (see Table S5 and Fig. S11) when compared with the parental mutant lines, This result was not surprising and was expected due to the stacking of mutations from different mutant sources, which impaired seed storage protein and starch accumulation, ultimately leading to reduced seed weight. Similar observations have been reported in barley bred for high lysine content [61]. However, this limitation could potentially be improved by crossing the progeny lines developed in the present study with elite wheat cultivars. This is a future objective of the study and has previously been demonstrated in high-lysine barley and quality protein maize [62–63].

### 4.5 PCR-based markers for breeding reduced-immunogenicity wheat

A total of 30 DNA markers were employed in this study to identify different deletions (Table S2). However, these markers do not comprehensively cover all immunogenic gliadins, underscoring the need for additional marker development. To address this limitation, we designed four new sub-genome specific markers targeting regions previously lacking coverage. In combination with existing markers, these newly developed markers enabled the validation of deletions identified through both genomic and protein analyses. Moreover, they provide a valuable resource for characterizing reduced-immunogenicity wheat lines developed in this study and for screening wheat germplasm in gene banks worldwide. These markers will be particularly important for preventing contamination in released reduced-immunogenic wheat lines and for screening materials following the introgression of deletion mutants into adapted genetic backgrounds.

## Conclusions

The present study comprehensively characterized a diverse collection of mutant wheat germplasm with reduced gluten content, targeting gliadins and glutenins associated with the *Gli1*, *Gli2*, and *Glu3* loci. The genetic resources examined included: (i) the ω-gliadin-free winter wheat cultivar Wasko^ω−^; (ii) the Gli-A2 and Gli-D2 deficient line Pegaso^(6A-6D)α−^; (iii) NILs in the Pegaso background carrying deletions at the *Gli-A1/Glu-A3*, *Gli-B1/Glu-B3*, *Gli-D1/Glu-D3*, *Gli-A1/Gli-D1*, and *Gli-B1/Gli-D1* loci; (iv) gamma-radiation mutants in the Jagger background with deletions at the *Gli-A1, Gli-B1 and Gli-D1* locus; and (v) the land-race San Pastor carrying a deletion at the *Gli-D1* locus. In addition, wheat cytogenetic stocks, including nullisomic-tetrasomic lines and CS deletion lines for group 1 and group 6 chromosomes, were evaluated. Although protein profiles for most of these mutants had been reported previously, their molecular characterization was conducted here for the first time using GlutEnSeq and molecular marker analyses. The integration of genetic, genomic, and proteomic approaches enabled the establishment of robust gene-to-protein relationships for key gluten loci, providing insights that were previously incomplete or only partially understood. Furthermore, crosses among selected mutant resources were performed to combine multiple mutations within single genetic backgrounds. These efforts resulted in the development of reduced-immunogenicity wheat lines in which major immunogenic gluten proteins were eliminated while preserving the accumulation of HMS-GS, which are critical for end-use quality. In conclusion, this study generated valuable genetic resources, including F₂ and F₃ populations carrying stacked mutations at the *Gli-1* and *Gli-2* loci, as well as genomic resources in the form of diagnostic molecular markers. Together, these resources provide a strong foundation for future wheat improvement efforts and serve as important pre-breeding materials for the development of reduced-immunogenicity wheat cultivars aimed at benefiting individuals affected by wheat allergy and celiac disease.

## CRediT authorship contribution statement

**Gaganjeet**: Writing-original draft, investigation, data curation; **Gautam Saripalli**: Writing-original draft, data curation and analysis; **Zachary Jones**: Writing original draft; **Anmol Kajla**: Resources; **Vijay Tiwari**: Resources; **Jacek Waga**: Resources; **Sachin Rustgi**: Conceptualization, Supervision, Writing– review & editing, Project administration, Funding acquisition.

## Acknowledgments

This work was supported by the U.S. Department of Agriculture, National Institute of Food and Agriculture (USDA-NIFA) under Grant No. SC-2022-10280. The authors are grateful to Dr. Stefania Masci, Dr. Francesco Sestili, and Dr. Domenico Lafiandra of the Dipartimento di Agrobiologia e Agrochimica, Università degli Studi della Tuscia, Viterbo, Italy, for generously providing the seeds of the Pegaso NILs.

## Data availability

The raw sequencing reads for all 13 GlutEnSeq samples have been deposited in the NCBI Sequence Read Archive (SRA) and are available under BioProject accession number PRJNA1457223. All other data supporting the findings of this study are included in the main text or the Supplementary Informati

