## Supplementary Figures for "Molecular Characterization of Gene–Product Relationships in the Wheat Gluten Complex for Reduced-Immunogenicity Wheat Development"

### Slide 1
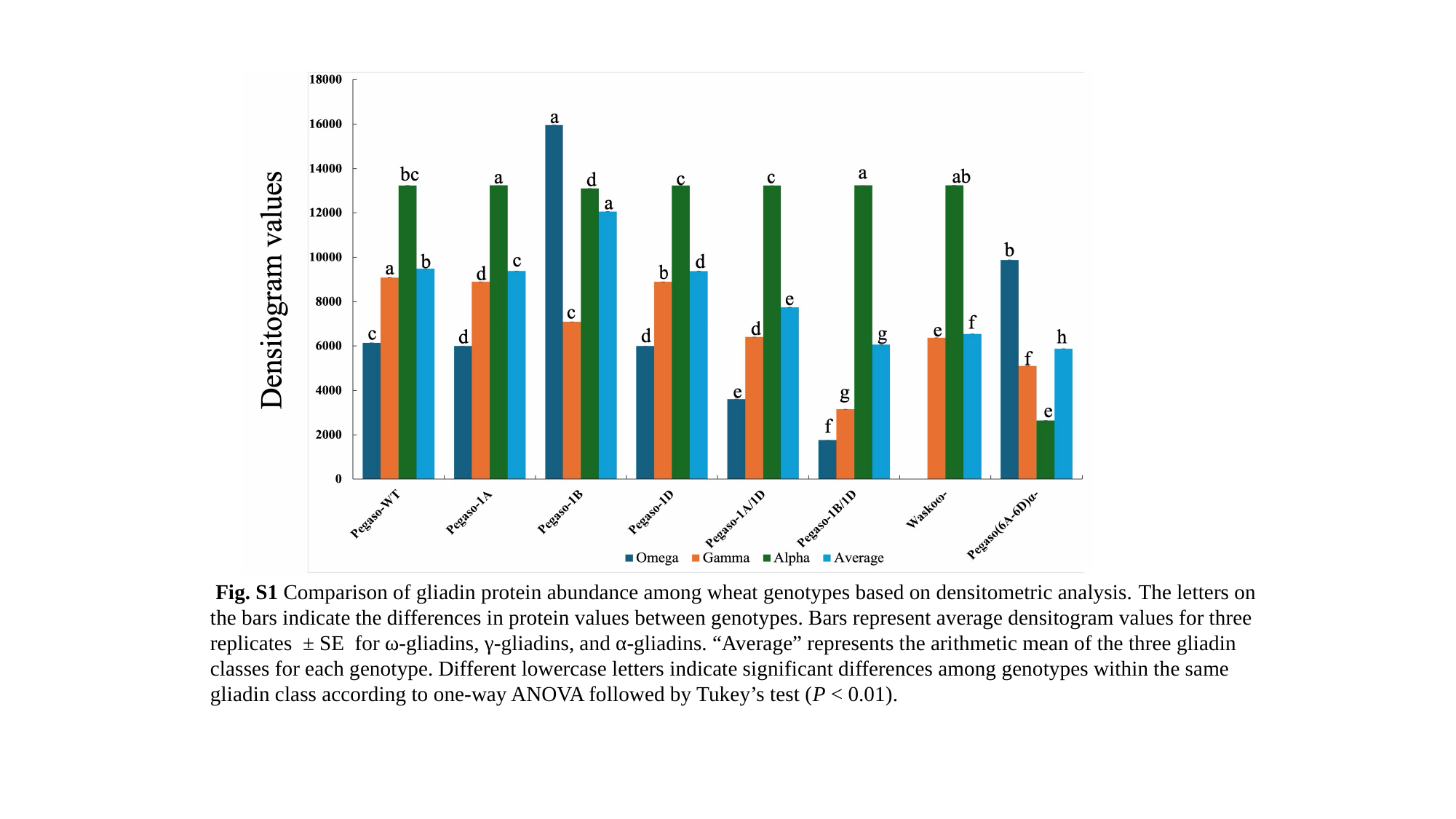

Fig. S1 Comparison of gliadin protein abundance among wheat genotypes based on densitometric analysis. The letters on the bars indicate the differences in protein values between genotypes. Bars represent average densitogram values for three replicates ± SE for ω-gliadins, γ-gliadins, and α-gliadins. “Average” represents the arithmetic mean of the three gliadin classes for each genotype. Different lowercase letters indicate significant differences among genotypes within the same gliadin class according to one-way ANOVA followed by Tukey’s test (P < 0.01).

### Slide 2
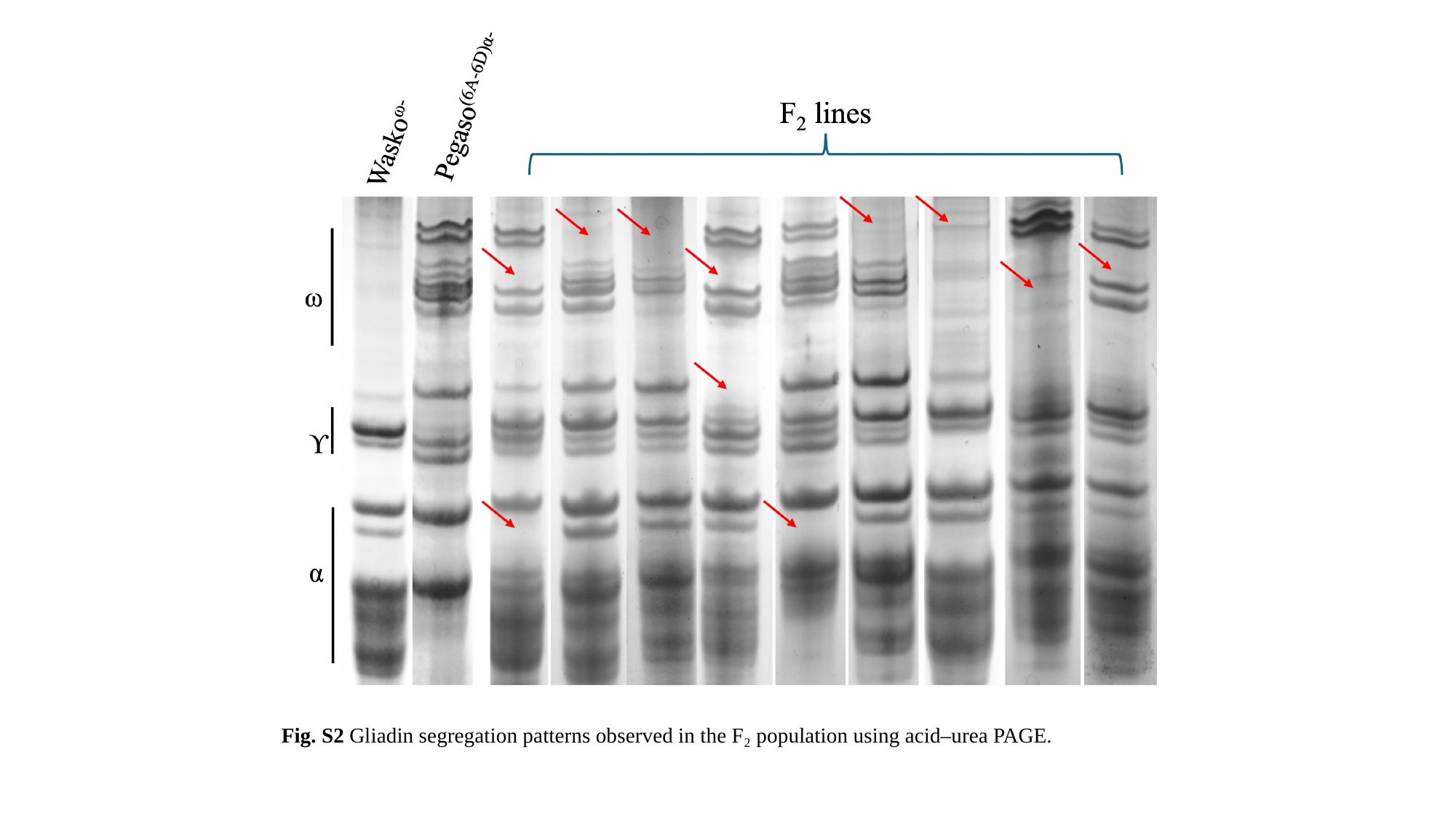

Fig. S2 Gliadin segregation patterns observed in the F₂ population using acid–urea PAGE.

### Slide 3
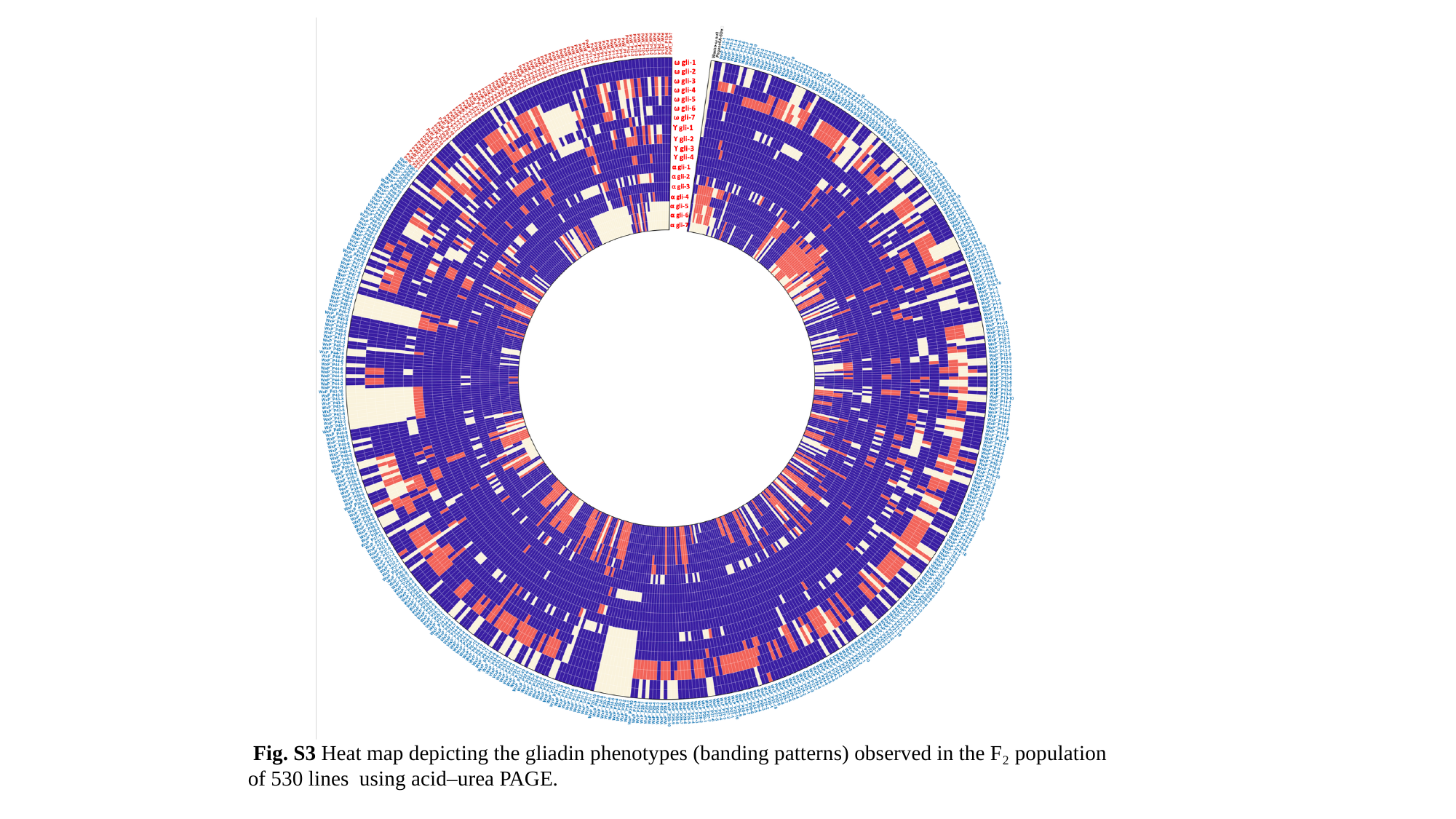

Fig. S3 Heat map depicting the gliadin phenotypes (banding patterns) observed in the F₂ population of 530 lines using acid–urea PAGE.

### Slide 4
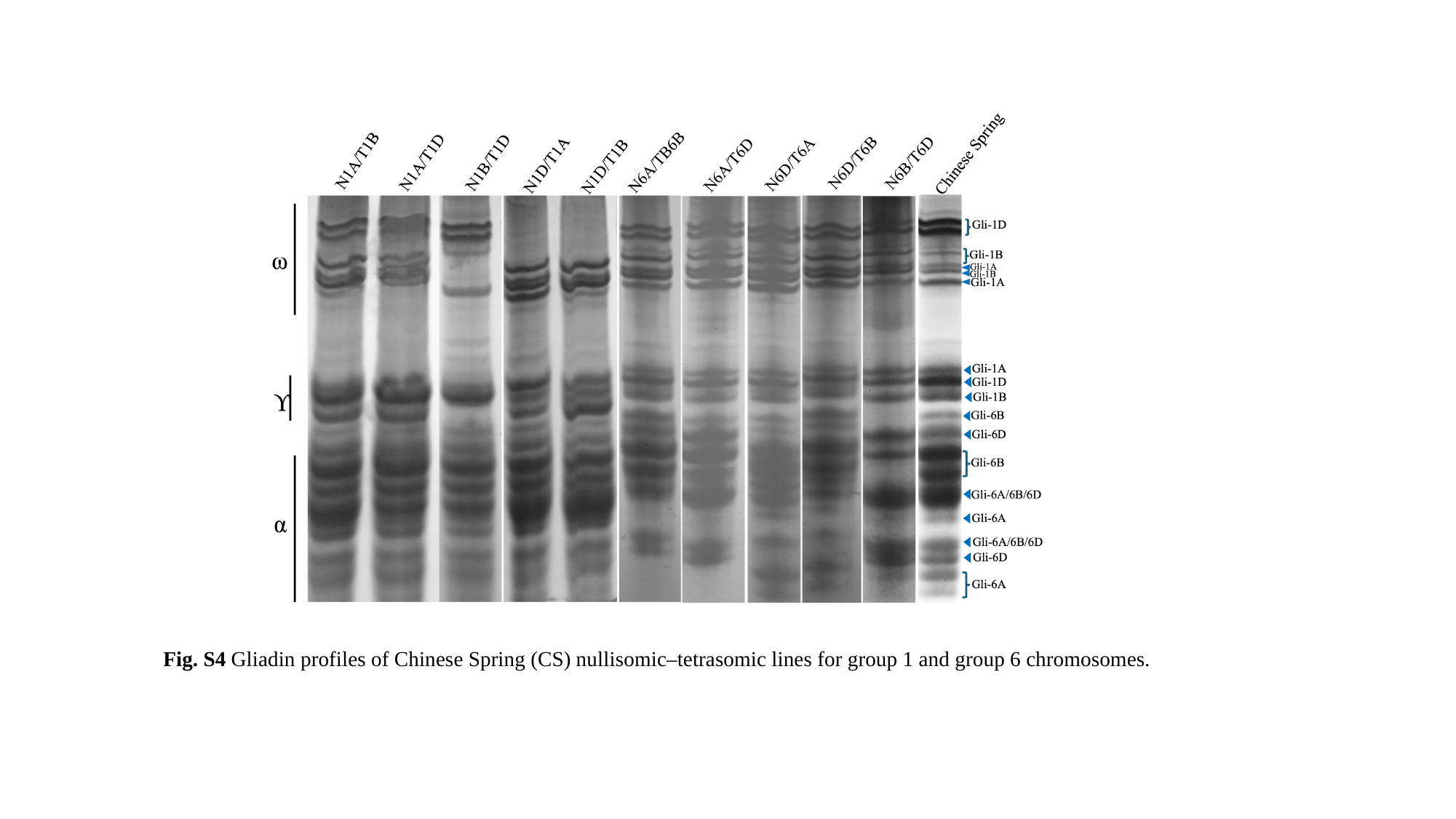

Fig. S4 Gliadin profiles of Chinese Spring (CS) nullisomic–tetrasomic lines for group 1 and group 6 chromosomes.

### Slide 5
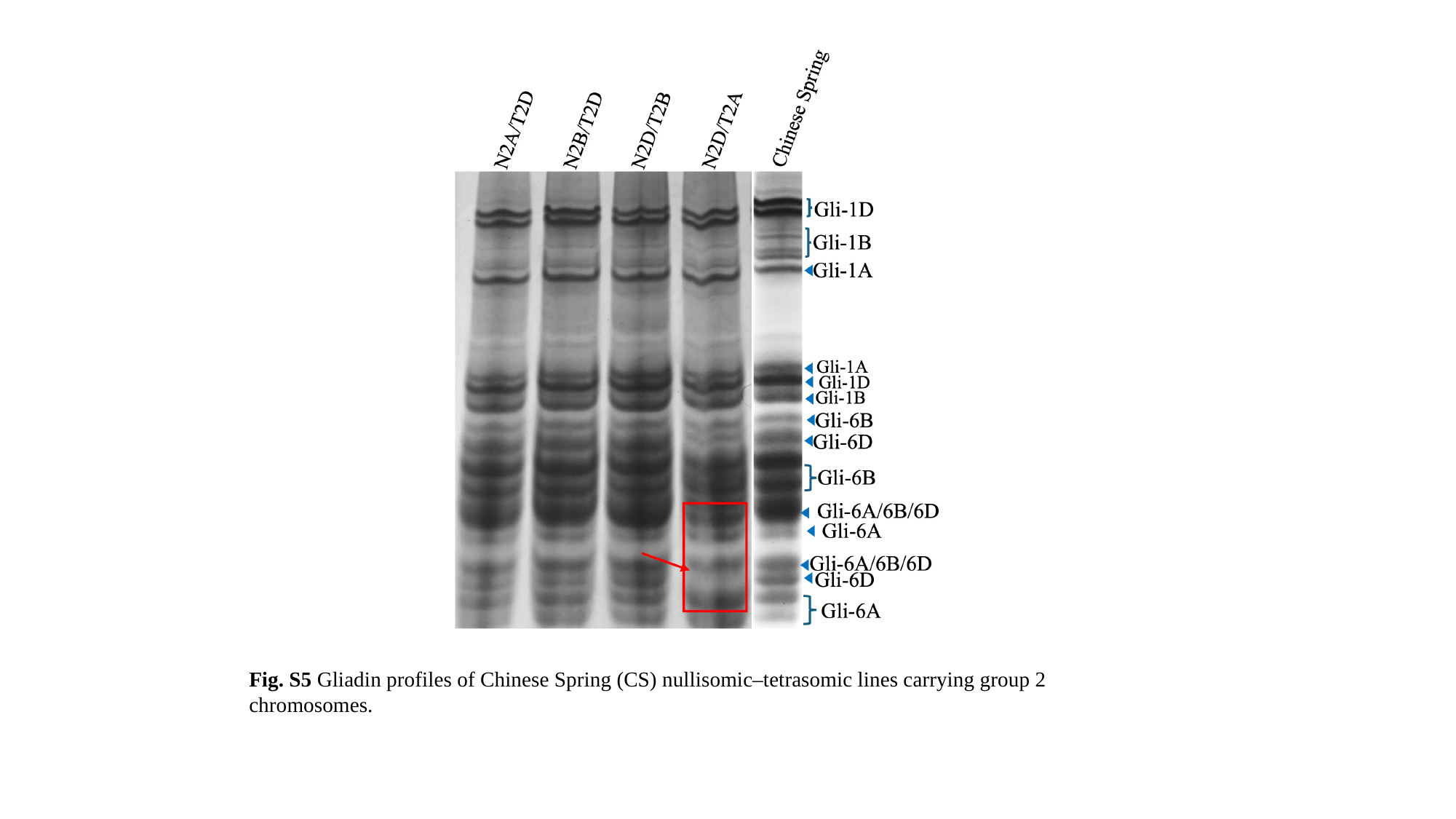

Fig. S5 Gliadin profiles of Chinese Spring (CS) nullisomic–tetrasomic lines carrying group 2 chromosomes.

### Slide 6
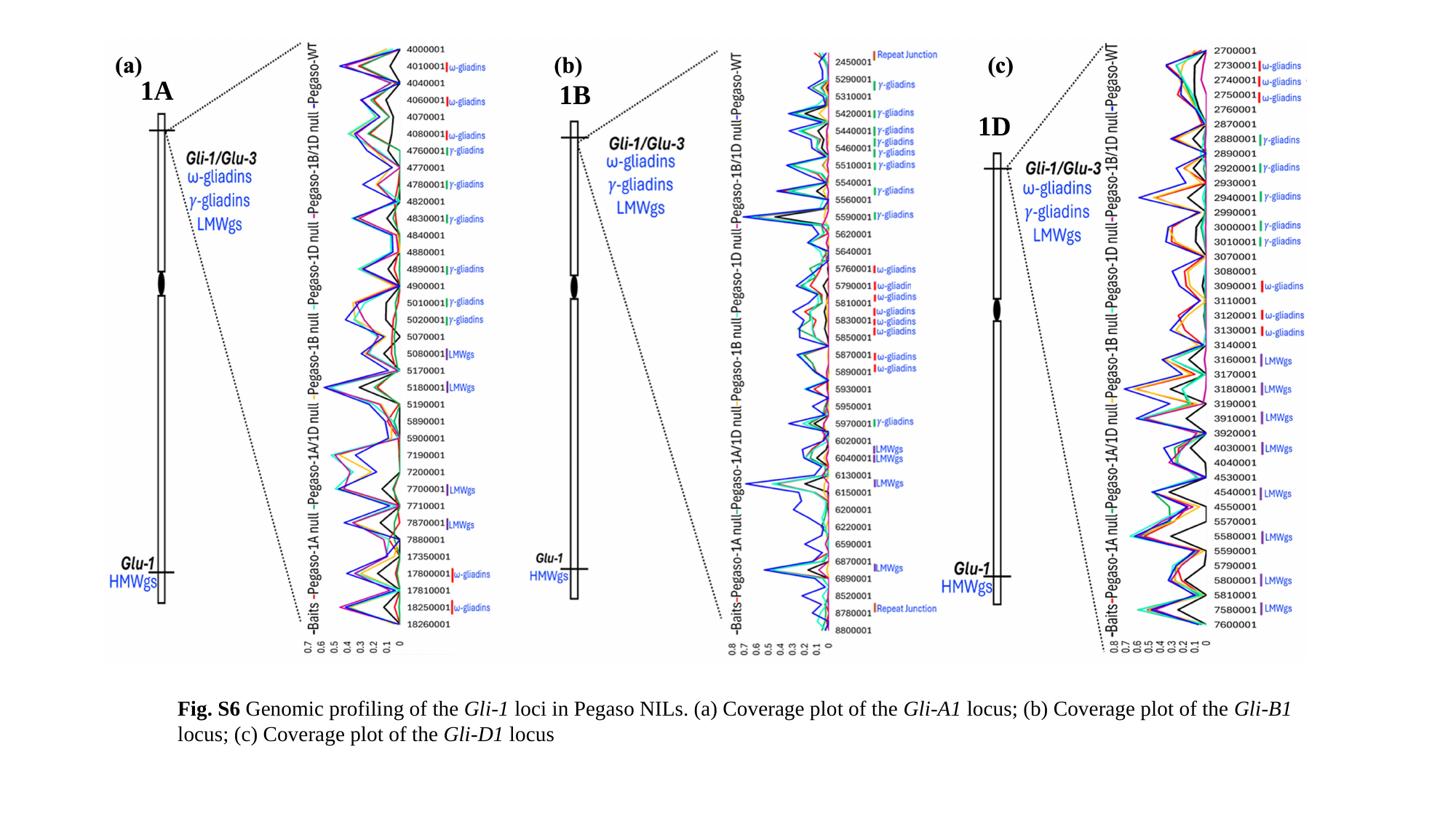

1A
1B
1D
Fig. S6 Genomic profiling of the Gli-1 loci in Pegaso NILs. (a) Coverage plot of the Gli-A1 locus; (b) Coverage plot of the Gli-B1 locus; (c) Coverage plot of the Gli-D1 locus

### Slide 7
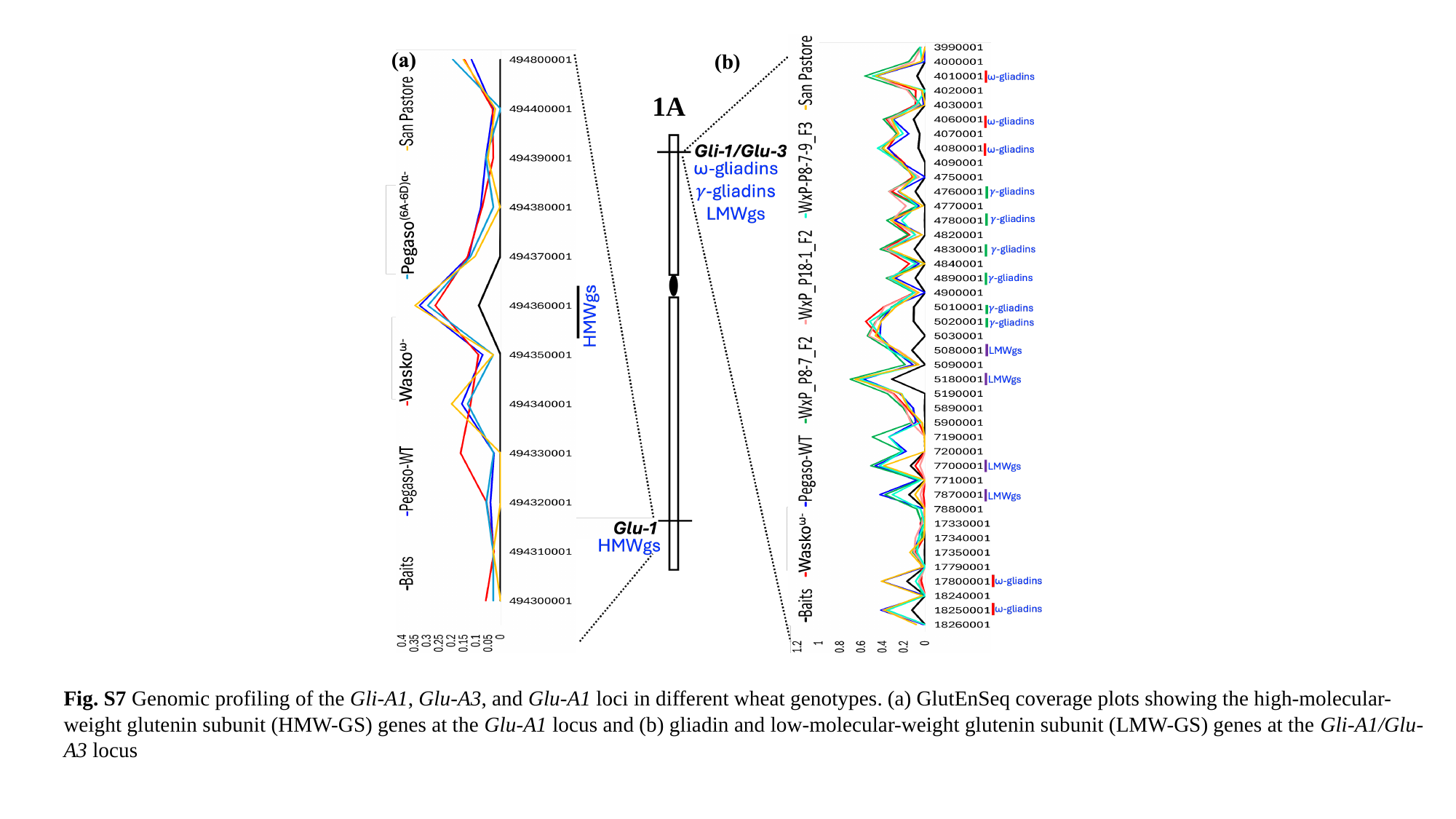

1A
Fig. S7 Genomic profiling of the Gli-A1, Glu-A3, and Glu-A1 loci in different wheat genotypes. (a) GlutEnSeq coverage plots showing the high-molecular-weight glutenin subunit (HMW-GS) genes at the Glu-A1 locus and (b) gliadin and low-molecular-weight glutenin subunit (LMW-GS) genes at the Gli-A1/Glu-A3 locus

### Slide 8
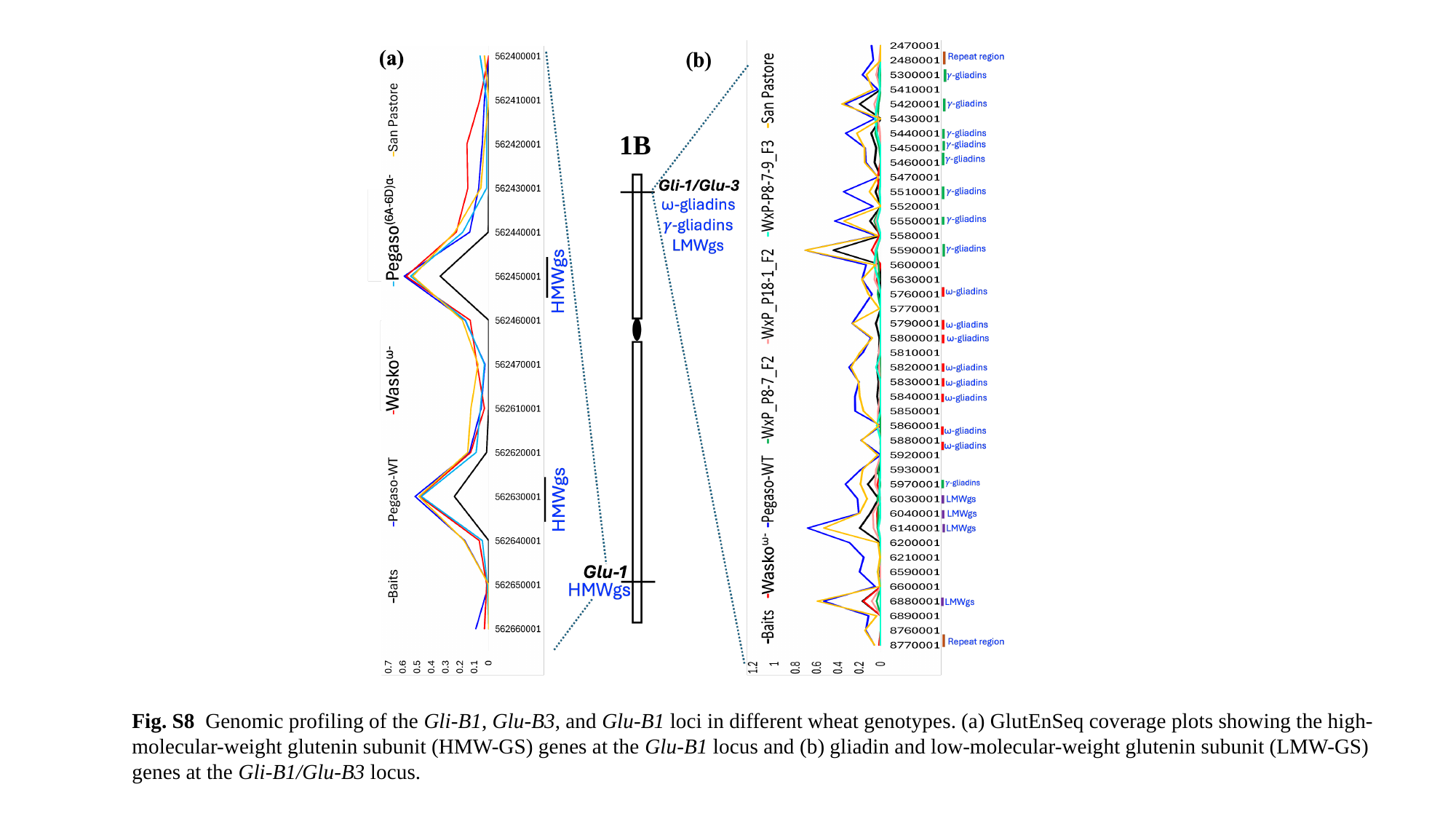

1B
Fig. S8 Genomic profiling of the Gli-B1, Glu-B3, and Glu-B1 loci in different wheat genotypes. (a) GlutEnSeq coverage plots showing the high-molecular-weight glutenin subunit (HMW-GS) genes at the Glu-B1 locus and (b) gliadin and low-molecular-weight glutenin subunit (LMW-GS) genes at the Gli-B1/Glu-B3 locus.

### Slide 9
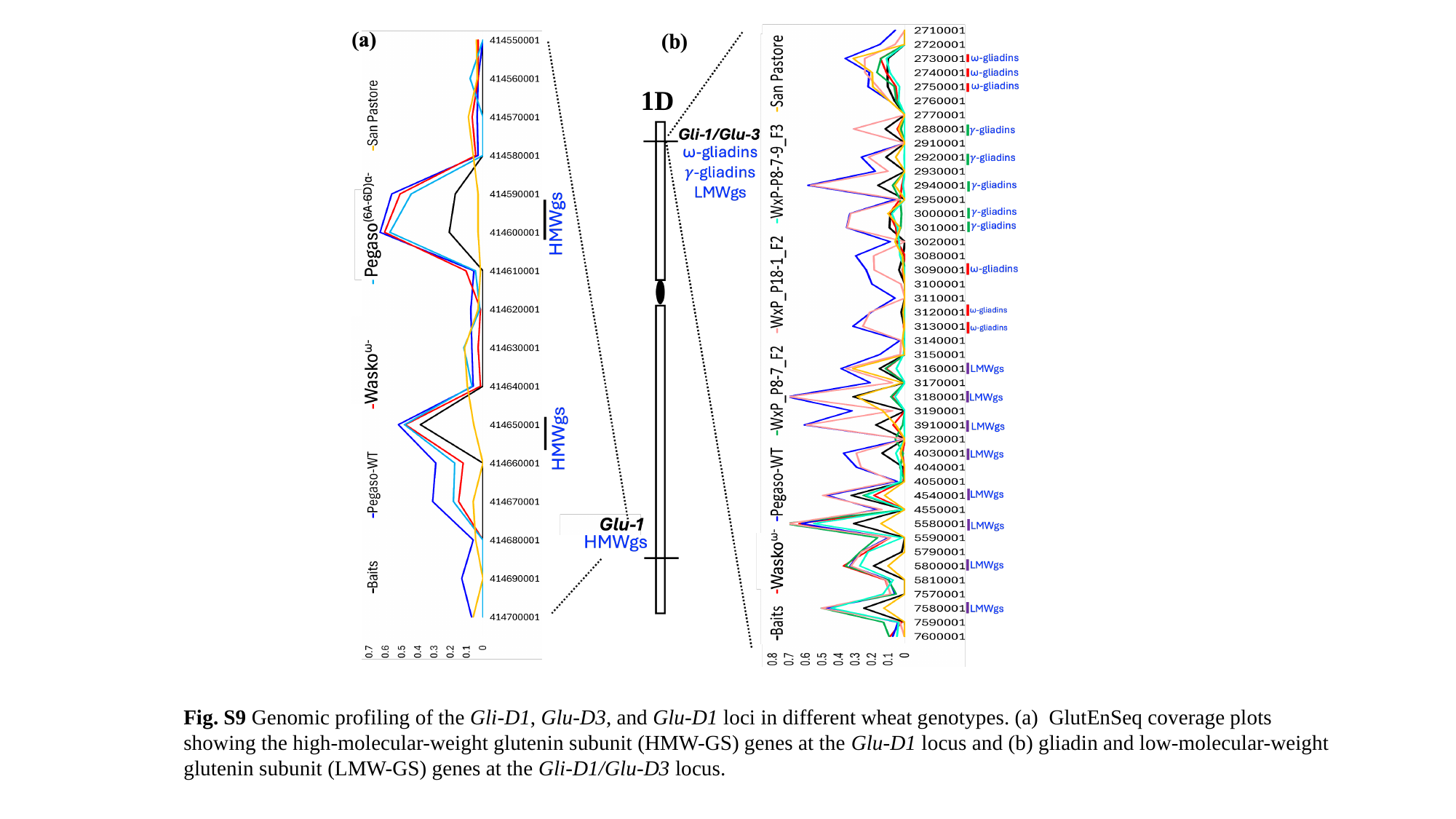

1D
Fig. S9 Genomic profiling of the Gli-D1, Glu-D3, and Glu-D1 loci in different wheat genotypes. (a) GlutEnSeq coverage plots showing the high-molecular-weight glutenin subunit (HMW-GS) genes at the Glu-D1 locus and (b) gliadin and low-molecular-weight glutenin subunit (LMW-GS) genes at the Gli-D1/Glu-D3 locus.

### Slide 10
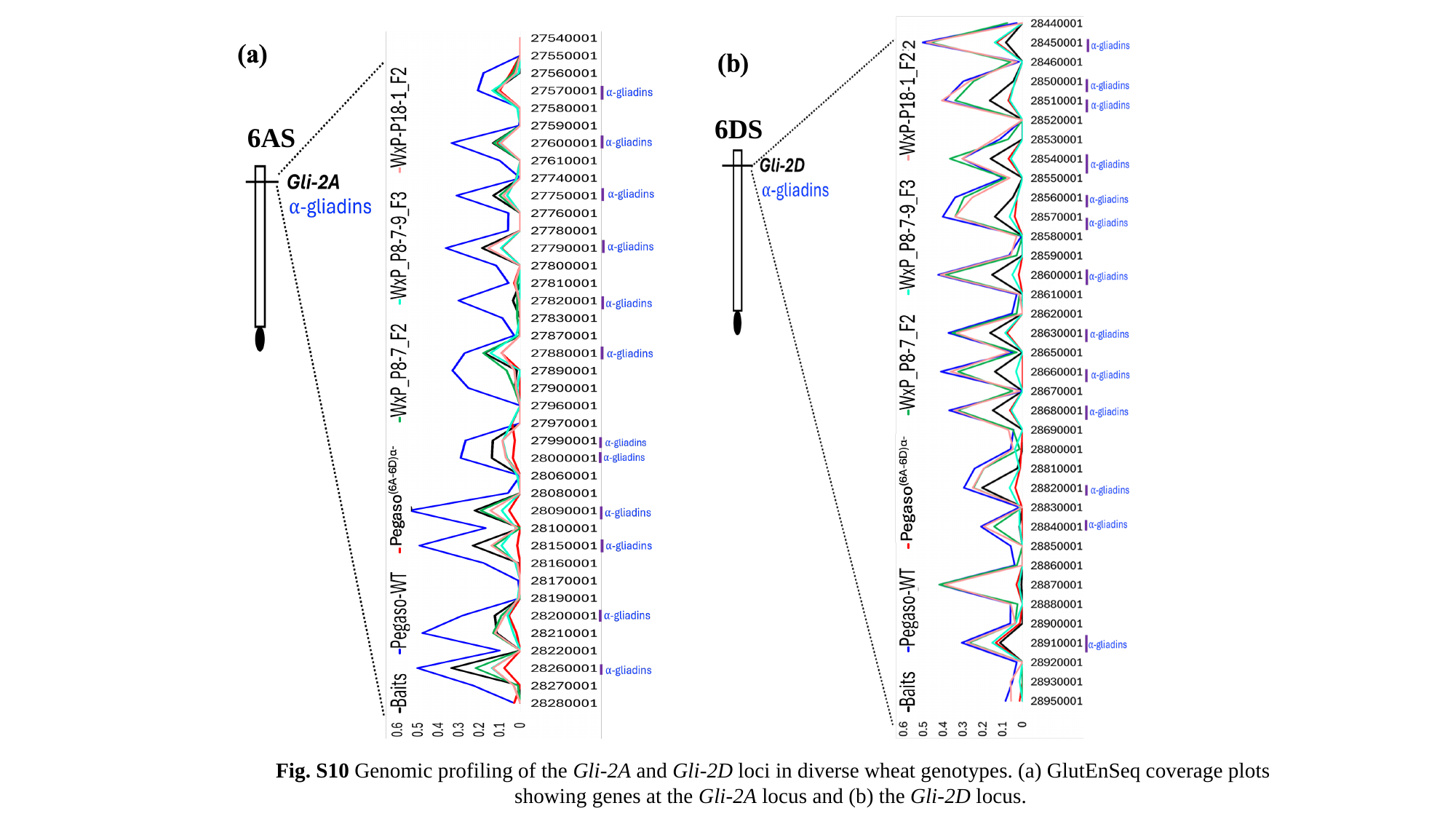

6DS
6AS
Fig. S10 Genomic profiling of the Gli-2A and Gli-2D loci in diverse wheat genotypes. (a) GlutEnSeq coverage plots showing genes at the Gli-2A locus and (b) the Gli-2D locus.

### Slide 11
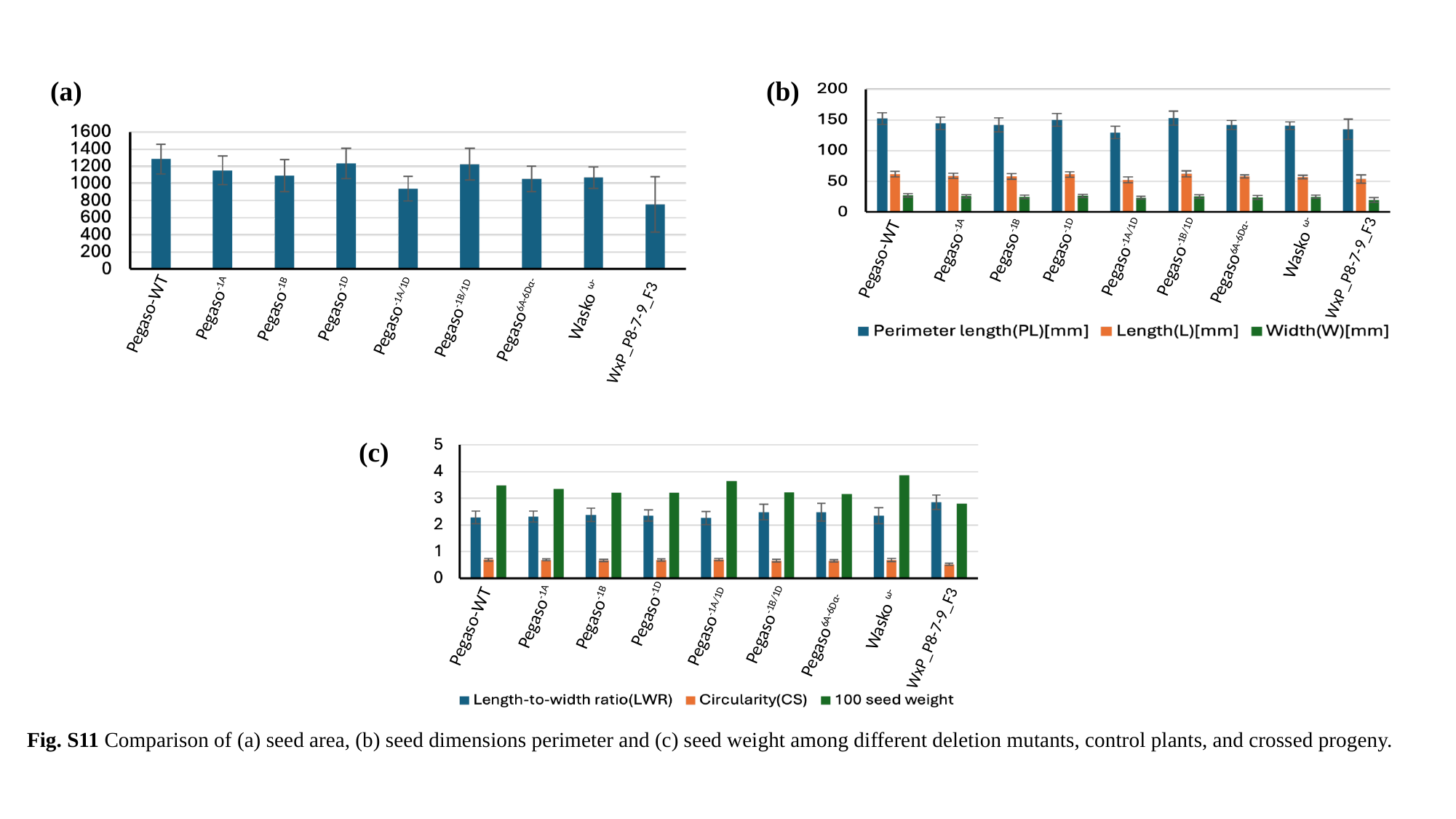

(a)
(b)
Wasko ^ω-^
 Pegaso^-1B^
 Pegaso^-1D^
 Pegaso^-1A^
 Pegaso^-1B/1D^
 Pegaso^-1A/1D^
Pegaso-WT
Pegaso^6A-6Dα-^
WxP_P8-7-9_F3
 Pegaso^-1D^
 Pegaso^-1A^
 Pegaso^-1B^
Pegaso^6A-6Dα-^
 Pegaso^-1A/1D^
Pegaso-WT
 Pegaso^-1B/1D^
Wasko ^ω-^
WxP_P8-7-9_F3
 Pegaso^-1D^
 Pegaso^-1B^
 Pegaso^-1B/1D^
Pegaso^6A-6Dα-^
 Pegaso^-1A/1D^
Wasko ^ω-^
Pegaso-WT
 Pegaso^-1A^
WxP_P8-7-9_F3
(c)
Fig. S11 Comparison of (a) seed area, (b) seed dimensions perimeter and (c) seed weight among different deletion mutants, control plants, and crossed progeny.

### Slide 12
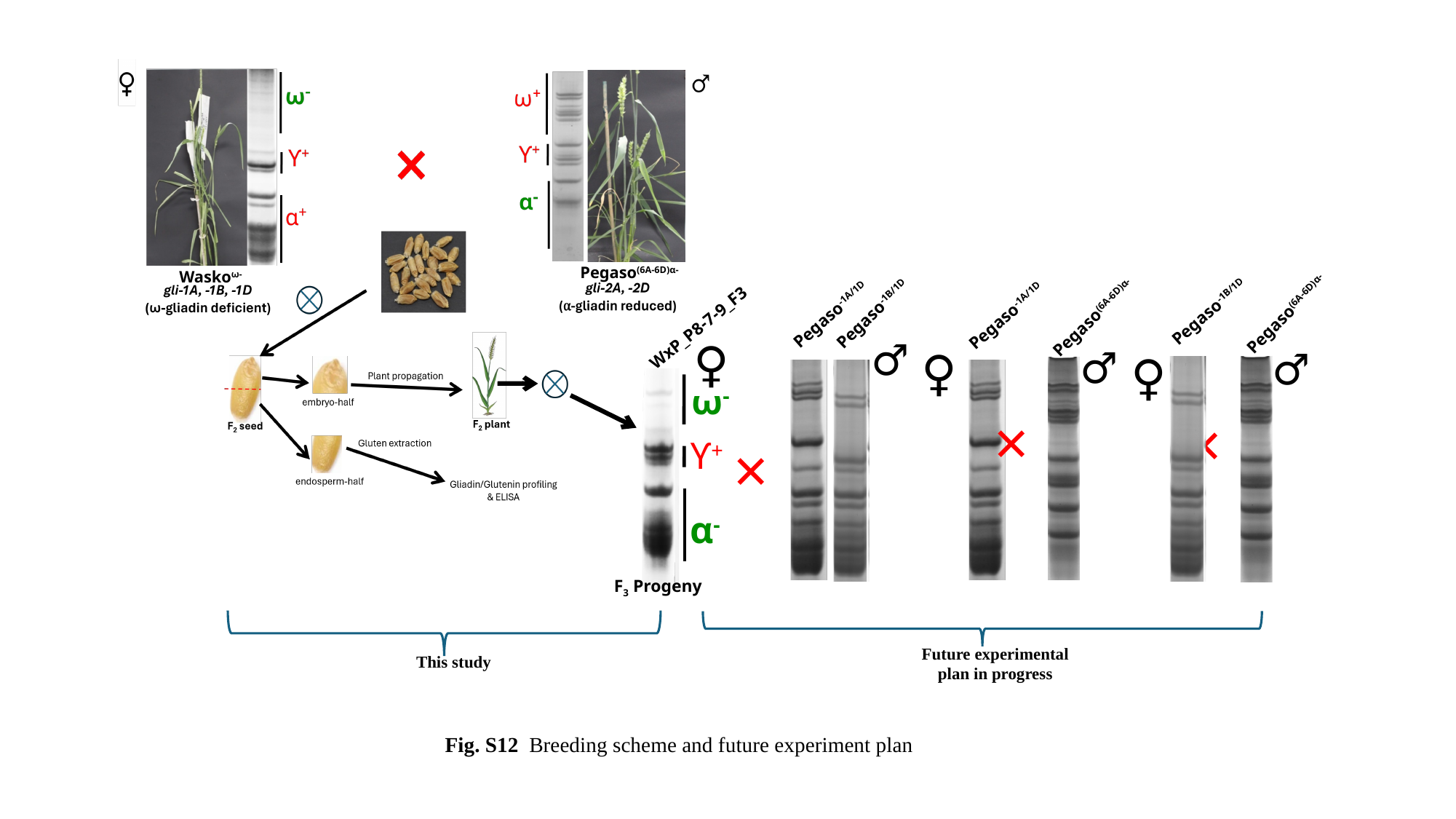

Pegaso^-1B/1D^
Pegaso^-1B/1D^
Pegaso^-1A/1D^
Pegaso^-1A/1D^
WxP_P8-7-9_F3
♀
♂
♂
♀
♂
♀
ω^-^
×
×
×
ϒ^+^
α^-^
F_3_ Progeny
Future experimental plan in progress
This study
Pegaso^(6A-6D)α-^
Wasko^ω-^
Pegaso^(6A-6D)α-^
Pegaso^(6A-6D)α-^
Fig. S12 Breeding scheme and future experiment plan
